# CyclinD1 controls development of cerebellar granule cell progenitors through phosphorylation and stabilization of ATOH1

**DOI:** 10.1101/2020.03.27.010926

**Authors:** Satoshi Miyashita, Tomoo Owa, Yusuke Seto, Mariko Yamashita, Shogo Aida, Tomoki Nishioka, Kozo Kaibuchi, Yoshiya Kawaguchi, Shinichiro Taya, Mikio Hoshino

**Affiliations:** Department of Biochemistry and Cellular Biology, National Institute of Neuroscience, NCNP, Tokyo 187-8502, Japan; Laboratory of Developmental Systems, Institute for Frontier Life and Medical Sciences, Kyoto University, Kyoto 606-8507, Japan; Department of Cell Pharmacology, Nagoya University Graduate School of Medicine, Nagoya 466-8550, Japan; Department of Clinical Application, Center for iPS cell Research and Application, Kyoto 606-8507, Japan

**Author notes:** Equal contribution (Satoshi Miyashita, Tomoo Owa). Correspondence: Department of Biochemistry and Cellular Biology, National Institute of Neuroscience, NCNP, 4-1-1 Ogawahigashimachi, Kodaira, Tokyo, 187-8502, Japan, E-mail.

**Keywords:** Development, Neural progenitor, Cerebellar granule cell, Proliferation, Differentiation, Atoh1, CyclinD1, Prox1, WNT signaling

## Abstract

Here we report that CyclinD1 (CCND1) directly regulates both the proliferative and immature states of cerebellar granule cell progenitors (GCPs). CCND1 not only accelerates cell cycle but also upregulates ATOH1 protein, an essential transcription factor that maintains GCPs in an immature state. In cooperation with CDK4, CCND1 directly phosphorylates Ser309 of ATOH1, which inhibits additional phosphorylation at S328, consequently preventing Ser328 phosphorylation-dependent ATOH1 degradation. PROX1 downregulates Ccnd1 expression by histone-deacetylation of Ccnd1 promoter in GCPs, leading to cell cycle exit and differentiation. WNT signaling upregulates PROX1 expression in GCPs. These findings suggest that WNT-PROX1-CCND1-ATOH1 signaling cascade cooperatively controls proliferation and immaturity of GCPs. We revealed that the expression and phosphorylation levels of these molecules dynamically change during cerebellar development, which was suggested to determine appropriate differentiation rates from GCPs to GCs at distinct developmental stages. This study contributes to understanding the regulatory mechanism of GCPs as well as neural progenitors.

## Introduction

During brain development, neural progenitors cease their proliferation and then differentiate into neurons. Neural progenitors have two characteristic states. They are initially proliferative and concurrently undifferentiated or in an immature state (Homem et al., 2015). At the appropriate timing, they stop proliferating and start differentiating into neurons, likely in response to extrinsic and intrinsic cues. Thus, regulation of proliferation and control of immaturity of neural progenitors are very crucial for proper brain development.

Cell cycle regulators such as cyclin family members and cyclin-dependent kinases (CDKs) promote the progression of cell cycle, while others including CDK inhibitors (CKIs) terminate the cell cycle in neural progenitors (Satyanarayana and Kaldis, 2009). For instance, combinatorial loss of D-type cyclins (Cyclins D1, D2 and D3), results in severe defects of cell proliferation during the brain development (Ciemerych et al., 2002). On the other hand, some basic helix-loop-helix (bHLH) transcription factors are known to either maintain neural progenitors in an immature state or make them differentiate into neurons (Dennis et al., 2019). For example, while overexpression of ATOH1, a bHLH protein, keeps cerebellar granule cell progenitors (GCPs) in immature state (Helms et al., 2001), NEUROD1 promotes differentiation of GCPs into the cerebellar excitatory neurons, granule cells (Butts et al., 2014b).

In the developing cerebral cortex, it has been suggested that there is a link between the immaturity of neural progenitors and their cell cycle length, especially G1 length (Salomoni and Calegari, 2010). CyclinD1 (CCND1) is well known to promote the transition from G1 to S phase in proliferating cells. Strikingly, the forced expression of CCND1 and its cofactor CDK4 keeps the apical progenitors, or radial glial cells, in the immature state, by prohibiting their transition into basal progenitors, while shortening their G1 length as expected from their classic function (Lange et al., 2009; Pilaz et al., 2009). Thus, it is believed that these two important steps, regulation of immaturity and control of proliferativity, are linked. However, the underlying molecular mechanisms remain unclear.

The cerebellum represents a good model system to address this question because of its well-described developmental processes and distinct layer structures that are composed of granule cell precursors (GCPs) and granule cells (GCs) (Roussel and Hatten, 2011). In mice, GCPs emerge from the rhombic lip as early as embryonic day 13.5 (E13.5) and migrate rostrally just beneath the pia mater to form the external granule cell layer (EGL). From E14.5 to two or three weeks after birth, they remain in an immature state and extensively proliferate in the outer EGL (oEGL), and subsequently differentiate into GCs while simultaneously exiting from the cell cycle in the inner EGL (iEGL). Finally, GCs migrate through the molecular layer and reach the inner granule cell layer (IGL) (Leto et al., 2016). Existence of an easy gene-transfer method, *in vivo* electroporation (Owa et al., 2018; Umeshima et al., 2007), for mouse GCPs also makes this model very useful.

Atonal homolog 1 (ATOH1), a bHLH transcription factor, is indispensable for maintaining GCPs in the immature state in the EGL. Loss of Atoh1 results in precocious differentiation from GCPs to GCs (Flora et al., 2009). In addition, ATOH1 has been suggested to have oncogenic potential in SHH-type medulloblastoma (Ayrault et al., 2010). Thus, precise regulation of ATOH1 protein levels is essential for proper development of GCPs and GCs. Levels of ATOH1 protein are known to be regulated by both transcriptional regulation of the Atoh1 gene and degradation of ATOH1 protein (Mulvaney and Dabdoub, 2012). As Atoh1 gene transcription is controlled by ATOH1 protein at the autoregulatory Atoh1 enhancer (Helms et al., 2000), degradation of ATOH1 protein decreases the autoregulatory transcription of this gene. The presence of this feedback regulation indicates that appropriate regulation of ATOH1 degradation is crucial for proper development of GCPs/GCs as well as prevention of tumorigenesis.

Studies have shown that phosphorylation of ATOH1 controls protein degradation of ATOH1 (Forget et al., 2014; Klisch et al., 2017; Zhao et al., 2008). For example, phosphorylation of serine (S) 328 of mouse ATOH1, which is recognized by E3 ubiquitin ligase Huwe1, is a trigger for proteasome degradation of ATOH1. Although several other studies have also identified the signaling pathways and/or molecules involved with ATOH1 degradation, the underlying machinery for ATOH1 degradation remains elusive. In this study, we classified GCPs into two subgroups; more proliferative/immature GCPs (AT+GCPs) that express ATOH1 and less proliferative/immature GCPs (ND+GCPs) that do not express ATOH1 but are NEUROD1-positive. The reduction of ATOH1 promotes GC development by accelerating transition from AT+GCPs to ND+GCPs (AT-ND transition), which eventually leads to differentiation into GCs. We found that CCND1 not only promotes cell cycle progression but maintains immaturity of GCPs by protecting against ATOH1 degradation. CCND1 and CDK4 cooperatively phosphorylate S309 of ATOH1, which suppresses further phosphorylation at S328, preventing ATOH1 degradation. Furthermore, the WNT-PROX1 pathway down-regulates CCND1 expression via chromatin histone deacetylation, which accelerates AT-ND transition of GCPs, eventually leading to GC development. We found that levels of the ATOH1 phosphorylation, CCND1, PROX1 proteins and WNT signaling vary during cerebellar development, which may control properly timed proliferation and differentiation of GCPs at distinct stages. These results give insights into the molecular machinery to coordinately regulate proliferation and immaturity in neural progenitors.

## Results

### ATOH1 expression controls the transition between subpopulations of GCPs

GCPs located in the oEGL have been reported to uniformly express ATOH1. However, recent studies revealed that there are some GCPs which do not express ATOH1. In order to characterize this population, we examined the developing cerebellum at postnatal day (P) 5 by immunostaining with ATOH1, NEUROD1 and KI67 (Fig. 1A). This revealed that while ATOH1-positive GCPs (AT+GCPs) were located in the superficial part of the EGL, some GCPs that were localized just beneath those AT+GCPs did not express ATOH1, as we and others have reported (Shiraishi et al., 2019; Xenaki et al., 2011). Interestingly, NEUROD1 was expressed in these ATOH1-negative GCPs (ND+GCPs) (Fig. 1B). Expression of ATOH1 and NEUROD1 was nearly mutually exclusive, indicating that ATOH1-negative GCPs and NEUROD1-negative GCPs in the EGL could be regarded as ND+GCPs and AT+GCPs, respectively. Our S-phase labeling with EdU incorporation for 30 min (Fig. S1A) and M-phase labeling with phospho-histone H3 (PH3) staining (Fig. S1B) confirmed that not only AT+GCPs but also ND+GCPs underwent DNA synthesis and mitosis in the EGL.

**Figure 1.**
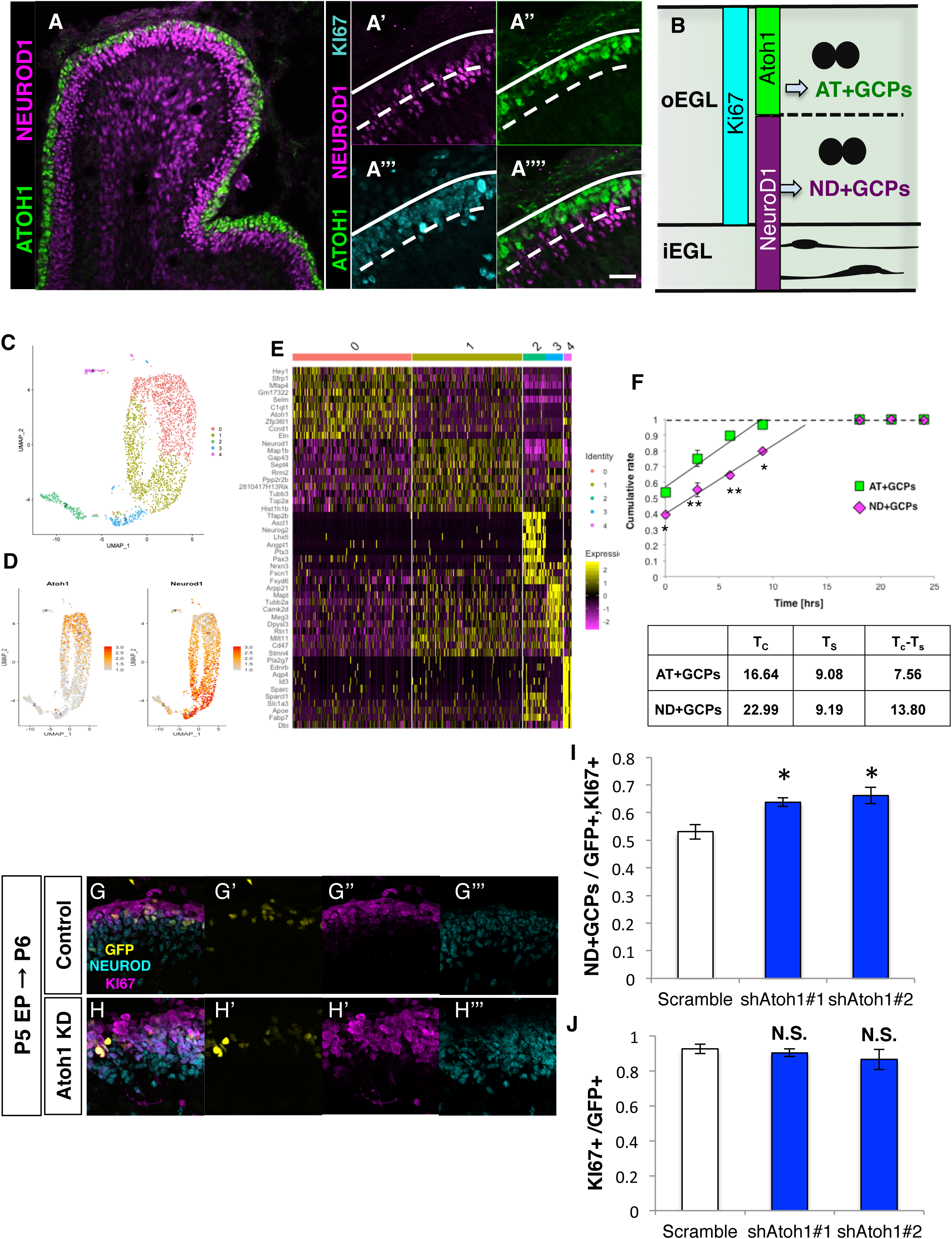
Characterization of AT+GCPs and ND+GCPs. Immunostaining of P5 cerebellum with indicated antibodies. (A) low magnification. (A’-A’’’) high magnification. (B) Schematic illustration of AT+GCPs and ND+GCPs. KI67-positive GCPs in the EGL were distinguished by the expression of two bHLH transcriptional factors, ATOH1 (AT+GCPs) and NEUROD1 (ND+GCPs). (C) *Pax6*-positive and *Pcna*-positive cells were extracted from published scRNA-seq data and analyzed with Seurat (see also the Experimental procedures). UMAP-based dimensional reduction and cluster analysis grouped extracted cells into 4 clusters. (D) Normalized expression of selected genes (*Atoh1*, *Neurod1*) were mapped onto the UMAP dimension. (E) Featured molecules in each cluster were determined and visualized by heatmap. Molecular features of each cluster revealed that clusters 0 and 1 were GCP lineage cells, cluster 3 was GCs and clusters 2 and 4 were likely to be interneuron precursors as *Pax3* and *Id3* expression were strong. In addition, the strong expression of *Atoh1* and *Neurod1* were observed in cluster 0 and cluster 1 respectively, suggesting that cluster 0 and cluster 1 might be AT+GCPs and ND+GCPs. (F) Cumulative labeling of AT+GCPs and ND+GCPs in P6 mice. Total cell cycle length (Tc) and S phase length (Ts) were calculated as previously described (see also Experimental Procedures). Calculated time of Tc, Ts and Tc-Ts were indicated in the table. The Student T-test was performed to test the difference of the ratio of EdU+ cells between AT+GCPs and ND+GCPs 0, 3, 6 and 9 hrs after EdU administration. N=3 for each time point. Two-way ANOVA analysis was performed against Cumulating rate from 0hrs to 9hrs in each cells, followed by Bonferroni test. (G-I) *Atoh1* KD analysis. (G, H) *Atoh1* shRNA was introduced to P5 mice EGL together with H2B-EGFP plasmid and immunostaining for NEUROD1 (Cyan) and KI67 (magenta) was performed 24 hrs later. Nuclei of electroporated cells were labeled with GFP (green). Images of Control (G) and *Atoh1* KD (H). (I) Ratios of NEUROD1+,KI67+ cells (ND+GCPs) among the electroporated GCPs (GFP+, KI67+ cells) were increased in the Atoh1 KD mice. (J) Ratios of KI67+ cells (total GCPs) were not significantly changed.. Student T-test was performed between Control (N=4) and *Atoh1* shRNA (sh#1: N=3, sh#2:N=3) electroporated mice. Mean± s.e.m. Scale bars: 20um(A, G, H).

Next, we utilized the published data of single-cell RNA sequencing (scRNA-seq) from P5 cerebella (Vladoiu et al., 2019). We bioinformatically extracted GCPs from all cerebellar cells with expression of *Pcna* and *Pax6* and clustered them, obtaining two groups containing GCPs (cluster 0 and cluster 1 in Fig. 1C). *Plxnb2*, *Barhl1*, *Mki67* and *Mcm6* expression confirmed that these clusters were GCPs (Fig. S1C). In addition, strong expression of *Atoh1* and *Neurod1* in cluster 0 and cluster 1 indicated that they corresponded to AT+GCPs and ND+GCPs, respectively (Fig 1D). We further generated a gene expression heatmap that showed that cluster 0 cells (AT+GCPs) and cluster 1 cells (ND+GCPs) have distinct gene expression profiles (Fig. 1E). While cluster 0 cells (AT+GCPs) tended to strongly express molecules involved with cell cycle progression or maintenance of undifferentiated states (such as *Sfrp1*, *Hey1* and *Ccnd1*), cluster 1 cells (ND+GCPs) strongly expressed molecules related to cell differentiation or migration (such as *Tubb3*, *Map1b* and *Gap43*) (Benito-Gonzalez and Doetzlhofer, 2014; Joesting et al., 2005). In addition, our cumulative labeling experiment to estimate cell cycle revealed that the total cell cycle length of ND+GCPs was significantly longer than that of AT+GCPs (Fig. 1F); the length of Tc-Ts, mainly affected by the length of G1, in ND+GCPs was twice as long as that of AT+GCPs. These findings suggest that AT+GCPs are more proliferative immature progenitors, while ND+GCPs are less proliferative and more differentiated progenitors. It was previously described that GCPs gradually move inward from the pial to the deeper side within the oEGL and then exit the cell cycle to become GCs in the iEGL (Chedotal, 2010). This suggested that within the oEGL, proliferative and immature AT+GCPs give rise to less proliferative and more differentiated ND+GCPs that eventually differentiate into postmitotic GCs in the iEGL.

By means of *in vivo* electroporation, we introduced short hairpin RNAs (shRNAs) for *Atoh1* (Atoh1 sh#1/ #2, Fig. S1D) into EGL cells facing the pia mater (AT+GCPs) at P5 and fixed brains 24 hours after electroporation. Nuclei of electroporated cells were visualized with co-injected H2B-GFP (Fig. 1G, H). In this experiment, the ratios of ND+GCPs in transfected cells were significantly increased in the Atoh1 knock-down (KD) cerebella compared to control (Fig. 1I). Although the ratio of GCPs (Ki67+ cells in the EGL) in electroporated cells was not affected by Atoh1 shRNAs one day after electroporation (Fig 1J), it was significantly reduced three days after electroporation (data not shown), consistent with the reported function of ATOH1 in GCP proliferation. Together, these observations suggest that ATOH1 is the key regulator that maintains GCPs at an immature and proliferative state and that reduction of ATOH1 leads to transition from AT+GCPs to ND+GCPs (AT-ND transition) that eventually accelerates differentiation into GCs.

### in vivo electroporation revealed Ccnd1 and Cdk4 are required for the AT-ND transition

Given the notion that ATOH1 reduction might be an important event for the “AT-ND transition” of GCPs, we tried to investigate its underlying machinery. We first focused on CCND1, because it was one of the featured genes for AT+GCPs (Fig. 1E) and targeted disruption of *Ccnd1* was reported to result in severe reduction of GCP proliferation (Pogoriler et al., 2006). From the scRNA-seq analysis of P5 cerebella, we confirmed that the expression of *Ccnd1* transcripts was much stronger in AT+GCPs (cluster 0) than ND+GCPs (cluster 1) (Fig. 2A). Immunostaining of P6 cerebella further showed that CCND1 protein expression was much stronger in AT+GCPs than in ND+GCPs (Fig. 2B). We re-clustered the scRNA-seq data of GCPs into three groups (G1, G2/M, S) by their “CELL CYCLE SCORE” (see experimental procedures). Interestingly, *Ccnd1* was detected in AT+GCPs at all cell cycle phases, while it was expressed in a cell cycle phase-dependent manner in ND+GCPs (Fig. S2A). Coimmunostaining of CCND1 and PH3 revealed the unexpected expression of CCND1 in the M-phase of AT+GCPs, while it was not observed in the M-phase of ND+GCPs similar to other dividing cells (Fig. S2B, C). These findings suggest that, in AT+GCPs, CCND1 may have functions other than promoting G1-S transition.

**Figure 2.**
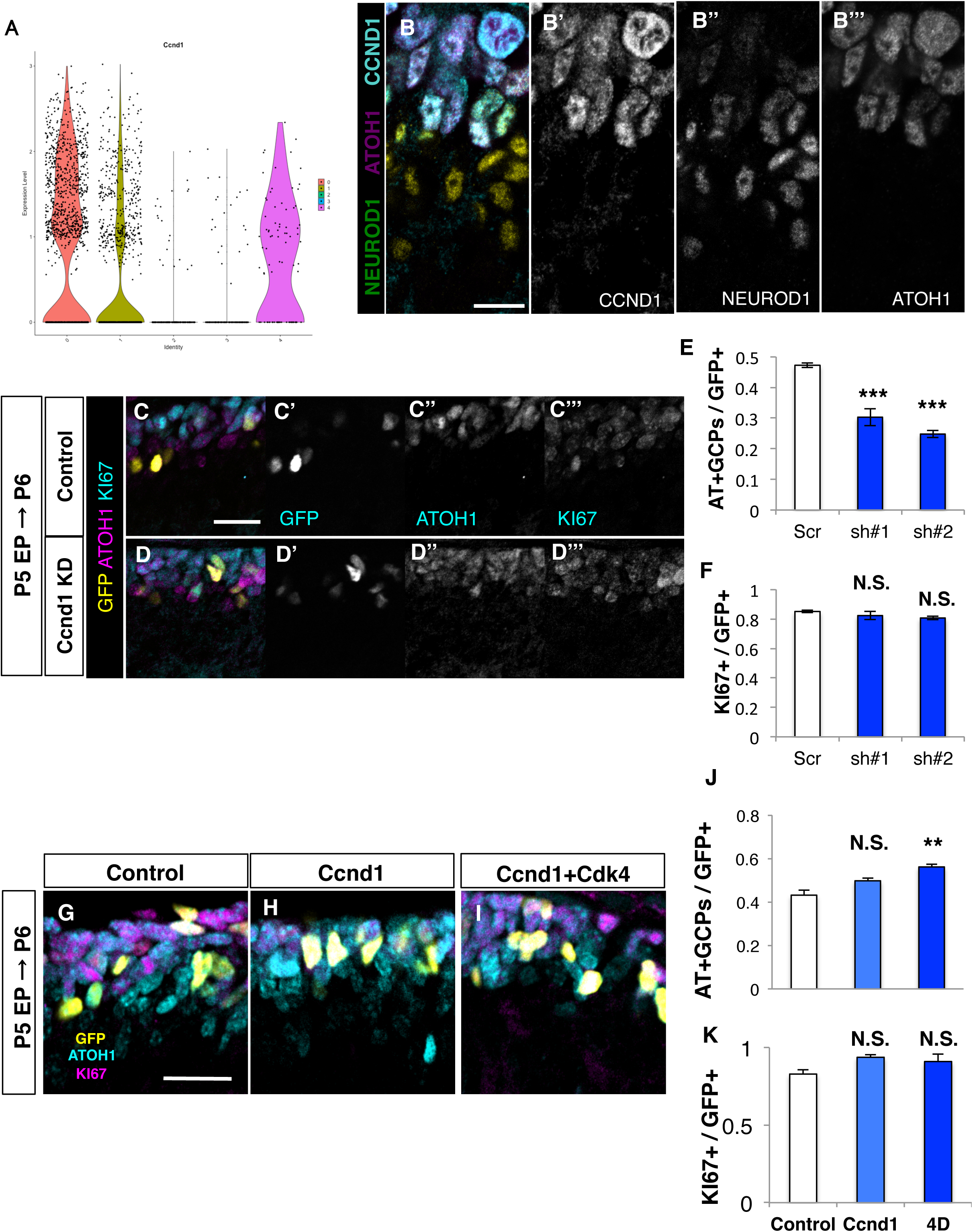
Requirement of CCND1 and CDK4 in the AT-ND transition. (A) Expression levels of *Ccnd1* in each cluster (Fig 1C) as indicated by violin plots. (B) Immunostaining of P5 cerebellum with indicated antibodies. (C-F) *Ccnd1* KD analysis. (C, D) *Ccnd1* shRNAs together with H2B-EGFP plasmid were introduced to P5 mice EGL and immunostaining for ATOH1 (magenta) and KI67 (Cyan) was performed the following day. Nuclei of electroporated cells were labeled with GFP (yellow). Images of Control (C) and *Ccnd1* KD (D). (E) Ratios of ATOH1+, KI67+ cells (AT+GCPs) among the electroporated cells was decreased in the *Ccnd1* KD mice. (F) Ratios of KI67+ cells (total GCPs) were not significantly changed. Student T-test was performed between Control (N=3) and *Ccnd1* KD (sh#1: N=3, sh#2:N=3) mice. Mean± s.e.m. (G-K) Over-expression analysis of *Ccnd1* with or without *Cdk4*. (G, H, I) *Ccnd1* and *Cdk4* expression plasmids were introduced to P5 mice EGL and immunostaining for ATOH1 (magenta) and KI67 (cyan) was performed the following day. Nuclei of electroporated cells were labeled with GFP (yellow). Images of Control (G), CCND1 over expression (H) and CCND1 and CDK4 over expression (I). (J) Ratios of ATOH1+, KI67+ cells (AT+GCPs) among the electroporated cells were not significantly changed between Control and CCND1. Forced expression of CCND1 and CDK4 significantly increased the ratio of AT+GCPs compared to Control. (K) Ratios of KI67+ cells (total GCPs) were not significantly changed. Tukey-Kramer test was performed on Control (N=6), CCND1 (N=4), CCND1 and CDK4 (N=3). Mean± s.e.m. Scale bars: 20um(B, C-D, G-I)

To investigate the role of CCND1 in GCPs, we performed KD experiment by introducing shRNAs for *Ccnd1* (*Ccnd1* sh#1/ #2, Fig. S2D) with H2B-GFP into AT+GCPs of P5 cerebella, which were fixed 24 hours after electroporation. Our immunostaining revealed that *Ccnd1* KD reduced AT+GCPs and increased ND+GCPs without affecting total GCPs (Ki67-positive cells) (Figure 2C-F and Fig S2E-G), suggesting that reduction of CCND1 promoted AT-ND transition. Next, we introduced expression vectors for CCND1 and CDK4 with H2B-GFP into P5 cerebella, which were fixed at P6. Although introduction of CCND1 alone did not affect the AT+GCP rate, simultaneous overexpression of CCND1 and CDK4 significantly increased AT+GCP population (Fig. 2G-K). This suggests that CCND1 may suppress AT-ND transition via the kinase activity of the CCND1/CDK4 complex rather than via its CDK-independent function.

### CCND1/CDK4 stabilizes ATOH1 protein through the direct phosphorylation

ATOH1 is a fragile protein that is rapidly degraded *in vitro*. Accordingly, administration of cyclohexamide (CHX), an inhibitor of protein translation, allowed clear visualization of GST-tagged ATOH1 (ATOH1-GST) degradation in Neuro2a (N2a) cells (Fig. 3A, B). However, this protein reduction was not observed with co-expression of CCND1 and CDK4 (Fig. 3A, B). This protection of ATOH1 degradation by CCND1/CDK4, however, could be blocked by administration with Palbociclib (Palb), a CDK4/6 inhibitor (Fry et al., 2004). Moreover, further administration of MG132, an inhibitor for proteasome-dependent protein degradation canceled the effect of Palb (Fig. 3C). Together with previous reports that Cyclin-Cdk-mediated phosphorylation of ID2, an HLH transcriptional regulator (Hara et al., 1997), promoted stabilization of this protein, these findings raised the possibility that CCND1/CDK4 stabilizes ATOH1 through phosphorylating ATOH1 via the kinase activity of CDK4.

**Figure 3.**
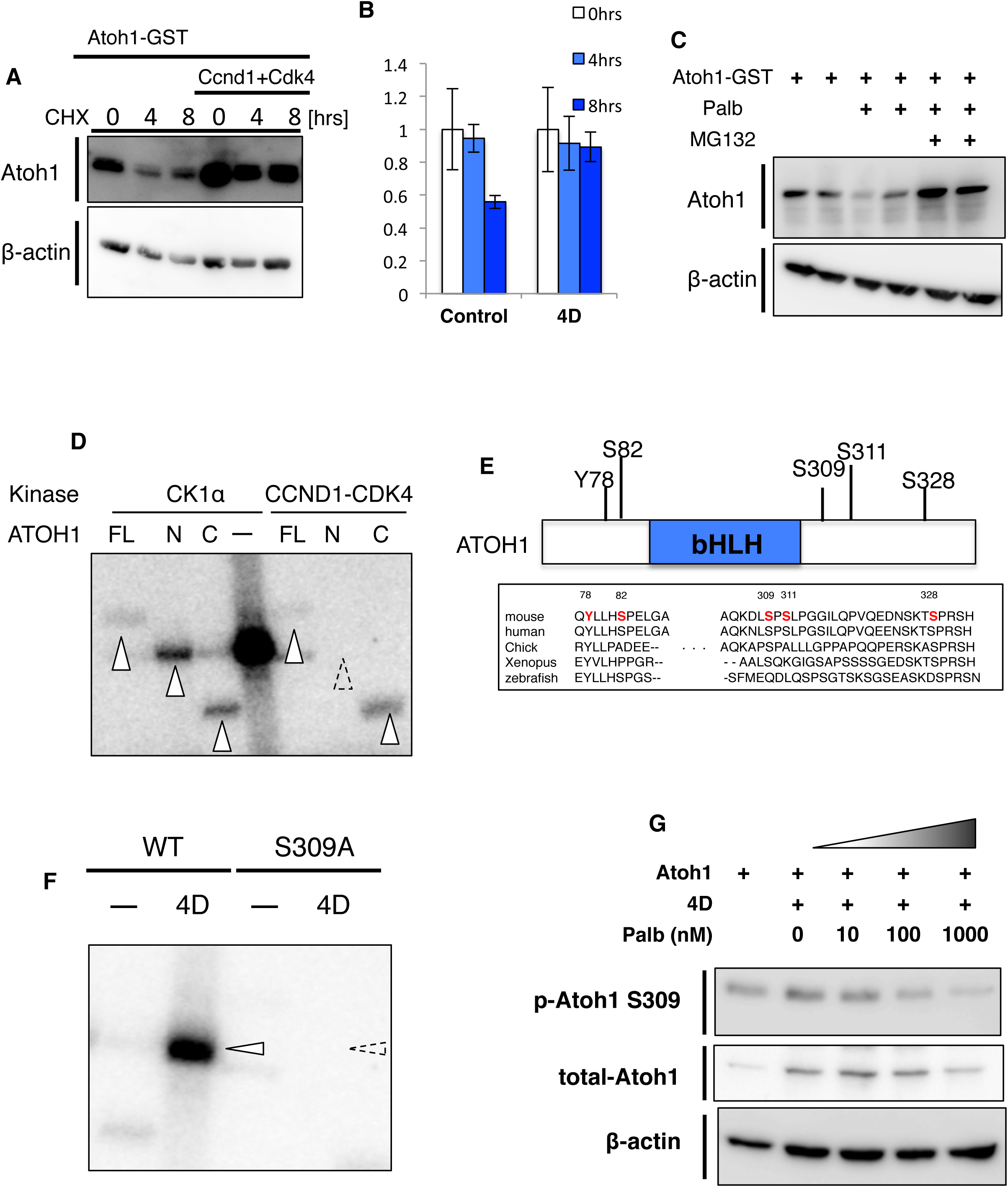
CCND1/CDK4 phosphorylates and stabilizes ATOH1 protein. (A, B) Cycloheximide (CHX) assay. (A) N2a cells transfected with Atoh1-GST, Ccnd1 and Cdk4 plasmids were harvested after 0, 4 and 8 hr treatments of 50ug/mL CHX and lysates were subjected to immunoblotting with anti-Atoh1 or β-actin as a housekeeping control. (B) Quantification of relative levels of ATOH1 to the β-ACTIN. Mean± s.e.m. from 3 trials of each condition. (C) N2a cells transfected with Atoh1-GST plasmids were treated with DMSO, Palbociclib (Palb) or Palb and 10 uM MG132 for 6hrs before harvest. Lysates of N2a were subjected to immunoblotting with anti-Atoh1 or β-actin. (D) in vitro phosphorylation assay of ATOH1. GST-tagged full length ATOH1 (FL), N-terminus ATOH1 including the bHLH domain (N) and C-terminus of ATOH1 downstream of the bHLH domain (C) were purified from N2a cells using glutathione-beads. ATOH1 proteins were incubated with either Casaine kinase 1 (CK1α) or CCND1 and CDK4 in the presence of [γ-32P]-ATP. Phosphorylation signals detected with FL, N and C by the phosphorylation of CK1α (lanes 1, 2, 3) are indicated by white arrowheads. Phosphorylation signals were also detected with FL and C by the phosphorylation of CCND1-CDK4 (lanes 5, 6, 7). Phosphorylation signals of N with CCND1-CDK4 were not detected. Strong CK1α self phosphorylation signals were observed (lane 4). (E) Phosphorylation sites of ATOH1-GST purified from N2a cells were determined by LC-MS/MS (see also Experimental Procedures). 5 phosphorylation sites (Y78, S82, S309, S311, S328) were identified, some of which are conserved across species. (F) In vitro kinase assay with either wild type ATOH1 (WT) or non-phosphorylated form of the Serine-to-Alanine 309 (S309A). WT or S309A were incubated with CCND1 and CDK4 (4D) in the presence of [γ-32P]-ATP. While WT with CCND1 and CDK4 gave rise to strong signals (lanes 1,2 white arrowheads), S309A showed no obvious signal (dashed arrow head indicated the predicted size of ATOH1). (G) Atoh1 expression plasmids were transfected into N2a cells with Ccnd1 and Cdk4 expression plasmids (4D). N2a cells were treated with 10 nM, 100 nM, 1000 nM Palb for 6 hrs before harvest. Lysates from N2a cells were subjected to immunoblotting with anti-Atoh1, phosphorylated ATOH1 S309 (p-Atoh1 S309) and β-actin as a housekeeping control.

It was previously suggested that CCND1 and CDK4 are involved in phosphorylation of ATOH1 in HEK293T cells (Tomic et al., 2018). However, it remained unclear whether CCND1/CDK4 directly phosphorylates ATOH1 or whether ATOH1 phosphorylation by CCND1/CDK4 participates in stabilization of ATOH1. Therefore, we performed *in vitro* phosphorylation assays with recombinant ATOH1, CDK4 and CCND1 proteins and isotope-labeled ATP (Fig. 3D). GST-tagged Atoh1, Ccnd1 and Cdk4 were transfected into HEK cells, respectively, and then purified with glutathione-conjugated protein sepharose beads. Casein kinase 1α (CK1α) was used as a positive control that is known to directly phosphorylate ATOH1 (Cheng et al., 2016)(Fig. 3D, lanes 1-3). A significant band was detected when CDK4 and CCND1 were co-incubated with full length (FL) ATOH1 (lane 5). We also performed this experiment for N-terminal and C-terminal regions of ATOH1 (N-ATOH1 and C-ATOH1), obtaining a strong signal for C-ATOH1 but not for N-ATOH1 (lanes 6, 7). These observations suggest that CCND1/CDK4 directly phosphorylate ATOH1, especially at the C-terminal region.

Some proteomics studies in the HEK293T cell line have identified several phosphorylation sites of ATOH1 protein (Cheng et al., 2016; Forget et al., 2014). Using ATOH1-transfected N2a cells, which possess more neuron-like features than HEK293T, we investigated the ATOH1 phosphorylation sites with LS-MS/MS. We confirmed that both CCND1 and CDK4 are endogenously expressed in N2a cells (data not shown). In this experiment, we identified five phosphorylation sites in the ATOH1 protein (Figure 3E). CCND1/CDK4 is known to phosphorylate proline-directed serine/threonine and our *in vitro* phosphorylation assay showed that CCND1/CDK4 directly phosphorylates the C-terminal region of ATOH1 (Fig. 3D). Although S309 and S328 are both highly conserved proline-directed serines in the C-terminal region (Fig. 3E), we excluded S328 as a candidate target for CCND1/CDK4 phosphorylation-mediated protein stabilization, because it was reported that phosphorylation of S328 leads to protein degradation of ATOH1. Therefore, we focused on S309 ATOH1. We performed *in vitro* phosphorylation assays using purified wild type (WT-ATOH1) and S309A-mutated (S309A-ATOH1) proteins. While WT-ATOH1 was robustly phosphorylated by CCND1/CDK4, S309A-ATOH1 was not phosphorylated (Fig. 3F). This suggests that CCND1/CDK4 directly phosphorylates ATOH1 at S309, at least, in vitro.

To further investigate phosphorylation at S309, we generated a specific antibody against S309-phosphorylated ATOH1 (p-S309-ATOH1). We first validated this antibody by immunoblotting with ATOH1 protein that was purified from N2a cells (Fig. S3A). As expected, this antibody recognized WT-ATOH1 but not S309A-ATOH1. However, when the purified WT-ATOH1 was de-phosphorylated by λ-phosphatase, no bands were detected. Moreover, no signals were detected with this antibody in N2a cell lysates transfected with S309A-ATOH1, while WT-ATOH1 gave rise to strong signals (Fig. S3B). These findings confirmed that our p-S309-ATOH1 antibody specifically recognized phosphorylation at S309 of ATOH1. In addition, we observed gradual reduction of phosphorylated ATOH1 at S309 in N2a cells transfected with ATOH1, CCND1 and CDK4, when Palb was administered at increasing concentrations of 10, 100, and 1000 nM (Fig. 3G). These results suggest that phosphorylation at S309 of ATOH1 by CCDN1/CDK4 contributes to increase in ATOH1 protein.

### Phosphorylation of ATOH1 S309 inhibits further phosphorylation at S328 to protect ATOH1 from protein degradation

As shown in Fig. 3A, coexpression of CCND1 and CDK4 suppressed ATOH1 degradation in cultured N2a cells in the presence of CHX. However, this effect by CCND1/CDK4 was not observed for S309A-ATOH1 (Fig. 4A). Next, we designed an expression plasmid for an ATOH1-phosphomimic form at S309 (S309D-ATOH1). In immunoblotting of transfected N2a cells, we observed much higher levels of S309D-ATOH1 compared with WT-ATOH1 (Fig. 4B).

**Figure 4.**
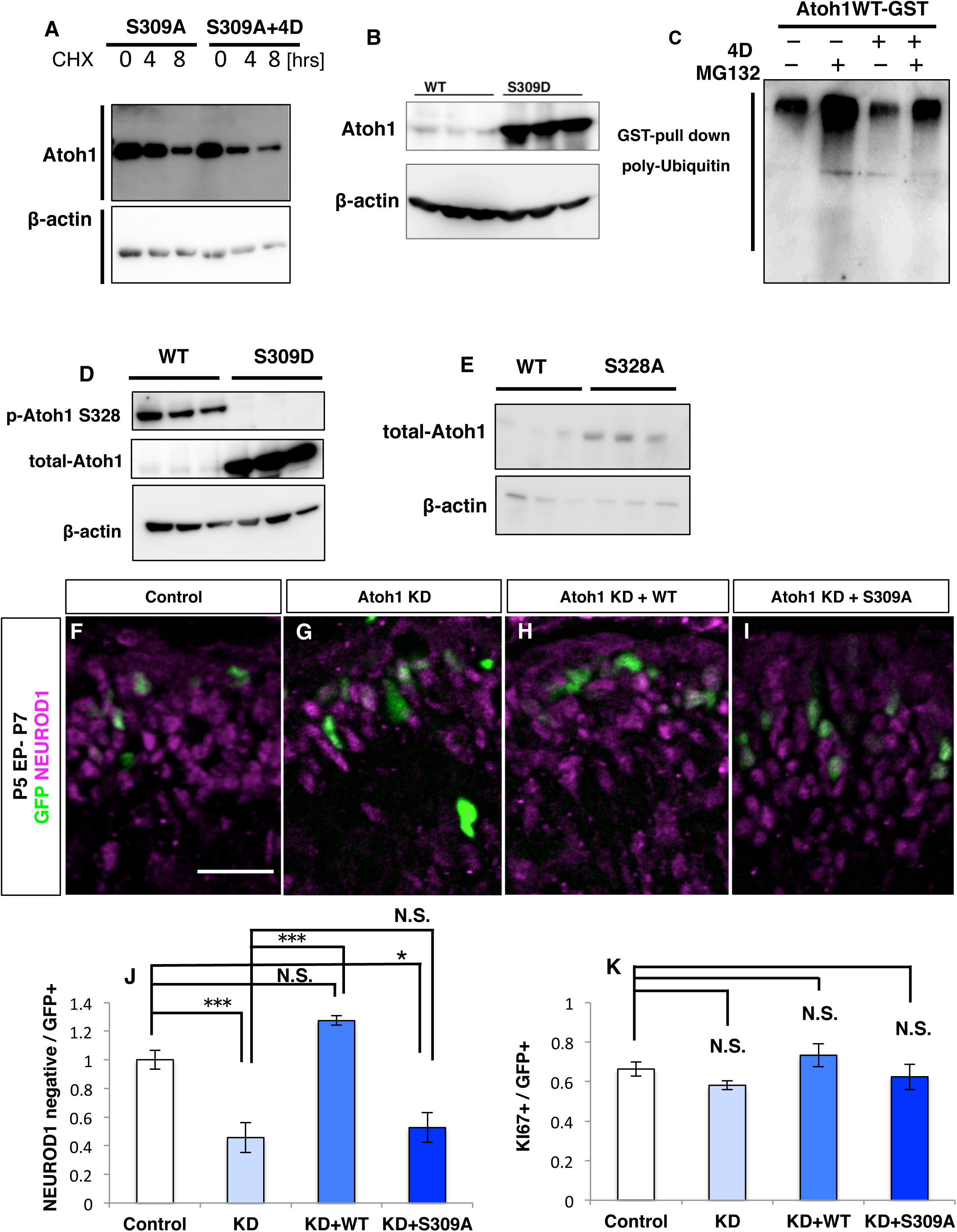
Phosphorylation of S309 inhibits further phosphorylation at S328A of ATOH1. (A) CHX assay with S309A. N2a cells transfected with either S309A or S309A+4D were treated with CHX for 0, 4 or 8 hrs before harvest. Lysates were subjected to immunoblotting with anti-Atoh1 or β-actin. (B) Lysates of N2a cells transfected with either WT or a phospho-mimic form of ATOH1 (S309D) were immunoblotted with anti-Atoh1 or β-actin. (C) GST-pull down assay of WT and WT with 4D. GST-Atoh1 plasmids were transfected with or without Ccnd1 and Cdk4 plasmids in N2a cells. Cell lysates were pull-downed with glutathione-conjugated beads after a 6 hr treatment of 10uM MG132 and then immunoblotted with anti-poly Ubiquitin. (D) Lysates of N2a cells transfected with either WT or S309D were immunoblotted to detect Atoh1 S328 phosphorylation (p-Atoh1 S328), Atoh1 or β-actin. (E) Lysates of N2a cells transfected with either WT or S328A were immunoblotted with anti-Atoh1 or β-actin. (F-K) Rescue experiment of Atoh1 KD with WT or S309A form of Atoh1. (F-I) Scramble (Control), *Atoh1* shRNA (Atoh1 KD), shRNA and WT (KD + WT) or shRNA and S309A (KD + S309A) plasmids were introduced to P5 mice EGL with H2B-EGFP plasmid and immunostaining for NEUROD1(magenta) was performed at P7. Nuclei of electroporated cells were detected by GFP (green). (J) Ratios of NEUROD1-, KI67+ cells (AT+GCPs) among the electroporated cells. (K) Ratios of KI67+ cells (total GCPs). Tukey-Kramer test was performed among the control (N=3), Atoh1 KD (N=3), Atoh1 KD +WT (N=3) and Atoh1 KD +S309A (N=3) mice. Mean± s.e.m. Scale bars: 20um(F-I)

We then investigated poly-ubiquitination of ATOH1 in N2a cells. ATOH1-GST was introduced to N2a cells, pull-downed with glutathione beads, and subjected to immunoblotting with an anti-poly-Ubiquitin antibody (Fig. 4C), which gave significant signals for poly-ubiquitination of ATOH1 (Fig. 4C, lane1). Addition of MG132 to the cell culture greatly increased the poly-ubiquitination (Fig. 4C, lane 2), confirming that ATOH1 is degraded via proteasome-dependent protein degradation. However, cotransfection of ATOH1-GST with CCND1 and CDK4 dramatically reduced polyubiquitination of ATOH1 (Fig. 4C, lane 3,4), indicating that CCND1/CDK4 suppresses poly-ubiquitination of ATOH1.

Phosphorylation of serine 328 (S328) of ATOH1 has been reported to lead to protein degradation mediated by the E3-ubiquitin ligase HUWE1. We generated an antibody against phosphorylated ATOH1 at S328 (p-S328-ATOH1) and performed immunoblotting to GST-WT-ATOH1 protein purified from transfected N2a cells. As expected, this antibody robustly detected purified GST-WT-ATOH1, but signals were lost upon λ-phosphatase coincubation (Fig. S4A). Additionally, this antibody detected strong bands in WT-ATOH1-transfected N2a cells but not in those transfected with S328A-ATOH1, a S328 phosphorylation-resistant form of ATOH1 (Fig. S4B). These observations suggest that our p-S328-ATOH1 antibody specifically recognizes phosphorylated ATOH1 at S328. We next transfected WT-ATOH1 or S309D-ATOH1 expressing plasmids into N2a cells, followed by immunoblotting with antibodies to p-ATOH1-S328 and total ATOH1. Interestingly, the phosphorylation signal at S328 of S309D-ATOH1 was dramatically decreased compared with WT-ATOH1, while the total amount of the protein was dramatically increased (Fig. 4D). Immunoblotting of transfected N2a cells found that the amount of S328A-ATOH1 protein was drastically increased compared with WT-ATOH1 (Fig. 4E), as was shown for HEK293T cells. These observations suggest that phosphorylation at S309 of ATOH1 by CCND1/CDK4 inhibits further phosphorylation at S328, which protects ATOH1 from degradation.

We further investigated the significance of phosphorylation at S309 ATOH1 during GC development *in vivo*. We electroporated WT-ATOH1 or S309A-ATOH1 to GCPs of P5 mice along with sh-Atoh1. Because the Atoh1 shRNA was designed to target the 3’ untranslated region (UTR) of Atoh1 mRNA, this KD vector effectively inhibits endogenous Atoh1 expression without affecting the expression of exogenous Atoh1 (Fig. S1D, E). KD of Atoh1 significantly decreased the proportion of AT+GCPs (NEUROD1-negative, KI67-positive cells) (Fig. 4J), as was observed in Fig. 1I. Although co-electroporation with WT-ATOH1 rescued the KD effect, co-introduction of S309A-ATOH1 did not have such a rescuing activity (Fig. 4F-K), implying that S309 phosphorylation of ATOH1 has a physiological importance in AT-ND transition of GCPs.

### PROX1 suppresses CCND1 expression through HDAC activity in GCPs

Previously, it was reported that PROX1 suppresses *Ccnd1* transcription in a neuroblastoma cell line(Foskolou et al., 2013). Although *Prox1* was reported to be expressed in the EGL (Lavado and Oliver, 2007), its precise localization within the EGL has not been shown. Our immunostaining detected strong expression of PROX1 in ND+GCPs and postmitotic GCs, and faint expression in AT+GCPs in the developing cerebellum (Fig. S5A-E). Double staining for CCND1 and PROX1 in the P6 cerebellum revealed complementary expression of CCND1 and PROX1, suggesting that *Ccnd1* may be transcriptionally regulated by PROX1 in the EGL (Fig. 5A).

**Figure 5.**
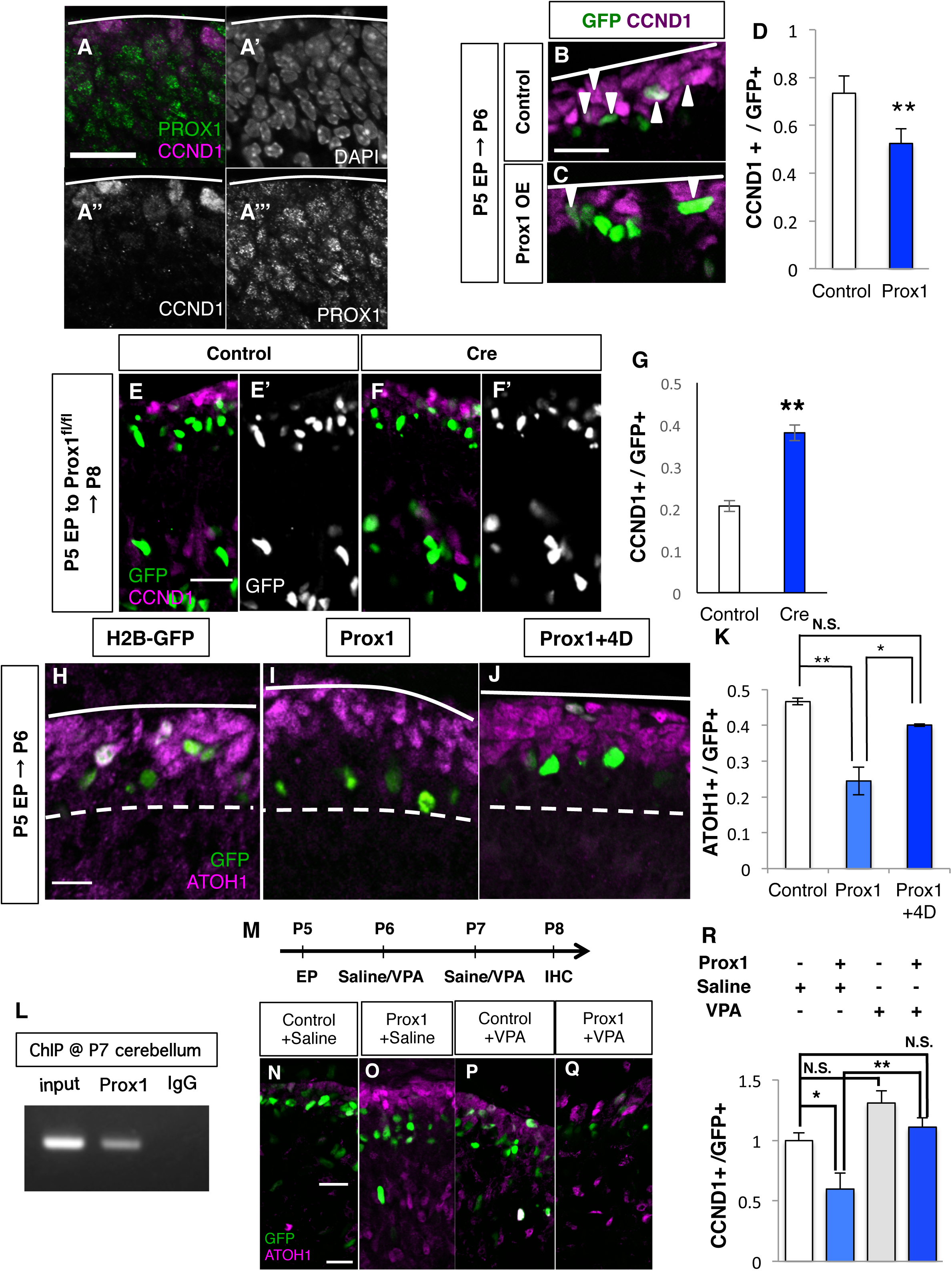
Prox1 suppresses CCND1 with HDAC activity. (A) Immunostaining of P5 cerebellum with indicated antibodies. (B-D) Overexpression analysis of PROX1. (B, C) Prox1 expression plasmids were introduced to mouse P5 EGL and immunostaining for CCND1 (magenta) was performed the following day. Nuclei of electroporated cells were detected with GFP (green). Images of Control (B), PROX1 overexpression (C). (D) The ratio of CCND1-positive cells among the electroporated cells. Student T-test between Control (N=3) and PROX1 overexpression (N=3). Mean±s.e.m. (E-G) *Prox1* acute KO analysis. (E, F) Expression plasmids of Cre recombinase were introduced to P5 EGL of *Prox1*^fl/fl^ mice and immunostaining for CCND1 (magenta) was performed at P8. Nuclei of electroporated cells were labeled with GFP (green). Images of H2B-GFP electroporation (E, Control), Cre electroporation (F, Cre). (G) The ratio of CCND1+, KI67+ cells among the electroporated cells. Student T-test between Control (N=3) and Cre (N=3). Error bar sem. (H-K) Rescue experiment of Prox1 with Ccnd1 and Cdk4-expressing plasmids. (H-J) Prox1 expressing plasmids were introduced with Ccnd1 and Cdk4 expression plasmids to P5 EGL together with H2B-GFP plasmid. Immunostaining for ATOH1 (magenta) was performed the following day. Nuclei of electroporated cells were detected by GFP (green). (K) Ratios of ATOH1+ cells (AT+GCPs) among the electroporated cells. The Tukey-Kramer test was performed on Control (N=6), Prox1 (N=3) and Prox1+4D (N=4) mice. Mean± s.e.m. (H) ChIP assay in P7 mouse cerebellum with anti-Prox1 or goat IgG as a negative control. (M-R) Administration of Valproic acid (VPA) to electroporated mice. (M) Expression plasmids of *Prox1* were introduced to P5 EGL and VPA or Saline was administered intraperitoneally 2 days after electroporation. Immunostaining for CCND1 (red) was performed the day following the last VPA treatment. Nuclei of electroporated cells were detected with GFP (green). (N-Q) Images of H2B-GFP with Saline (N), PROX1 with Saline (O), H2B-GFP with VPA (P) and PROX1 with VPA (Q). (R) The ratio of CCND1+ cells among the electroporated cells. Tukey-Kramer test among Control (N=4), PROX1 (N=4), Control +VPA (N=3) and PROX1 +VPA (N=4). Mean± s.e.m. Scale bars: 20um (A-B, E-F, H-J, N-Q)

When we introduced the PROX1 expression plasmid into P5 EGL, we observed that CCND1-expressing cells were significantly reduced 24 hours after electroporation (Fig. 5B-D). In contrast, introduction of a Cre recombinase-expressing vector to the EGL of homozygous Prox1-floxed (*Prox1*^fl/fl^) mice resulted in an increase in CCND1-positive cells (Fig. 5E-G). Together with a previous report that PROX1 suppresses expression of CCND1 in a neuroblastoma cell line, these findings suggest that PROX1 downregulates CCND1 expression in GCPs in the developing cerebellum. We also found that PROX1 overexpression reduces AT+GCPs (Fig. 5H, I, K) and increases ND+GCPs (Fig. S5F, G) without affecting the total GCP (Ki67-positive cells) number 24 hours after electroporation. However, these effects of PROX1 were canceled by co-expression of CCND1/CDK4 (Fig. 5J, K, data not shown). These observations suggest that PROX1 accelerates AT-ND transition by suppressing expression of CCND1 in GCPs.

A previous study demonstrated that PROX1 recruits HDACs to suppress expression of downstream genes through histone deacetylation at their promoter regions (Steffensen et al., 2004). We performed chromatin immunoprecipitation (ChIP) assay on the postnatal cerebellum using a Prox1 antibody and observed binding of PROX1 to the promoter region of *Ccnd1* (Fig. 5L). To evaluate the involvement of HDACs in suppressing CCDN1 expression by PROX1, we electroporated P5 mice with PROX1 and H2B-GFP and intraperitoneally injected VPA (an inhibitor for class I/II HDACs) at P6 and P7, followed by fixation at P8 (Fig. 5M). While PROX1 overexpression significantly reduced CCND1-positive cells, such an effect was clearly canceled by VPA administration (Fig. 5N-R). This suggests that HDACs may function under the control of PROX1 to suppress the CCND1 expression, which eventually leads to AT-ND transition in the EGL.

### Dynamic expression of CCND1, ATOH1 and PROX1 during cerebellar development controls the timing of the AT-ND transition

We and others have previously suggested that proliferation of GCPs and their differentiation to GCs are precisely regulated in a developmental stage-dependent manner (Miyashita et al., 2017): GCPs tend to proliferate at early stages, while they preferentially differentiate at later stages. To confirm this, cerebella were electroporated with H2B-GFP at P5 or P10, followed by fixation 24 hours after electroporation. Using electroporation, H2B-GFP was introduced to AT+GCPs located just beneath the pia mater. One day after electroporation, the ratio of AT+GCPs (or NEUROD1-negative cells in Fig. 6A) was larger at the earlier stage than at the later stage (Fig. 6A), suggesting that AT-ND transition occurs more frequently at later stages during the cerebellar development. This tendency was also observed in cultured GCPs. GCPs were purified from P5 or P10 cerebella and cultured for one or two days in vitro (DIV1 or DIV2). In our experimental conditions, more than 90% of GCPs (Ki67-positive cells) derived from either P5 or P10 cerebella were AT+GCPs (NeuroD1-negative Ki67-positive cells) at DIV1 (Fig. 6B-F). However, at DIV2, the rate was reduced to 70% for GCPs derived from P10 cerebella, but unchanged for those from P5 cerebella. This also confirmed that GCPs derived from the later stage cerebellum elicit AT-ND transition more frequently than those at the earlier stage.

**Figure 6.**
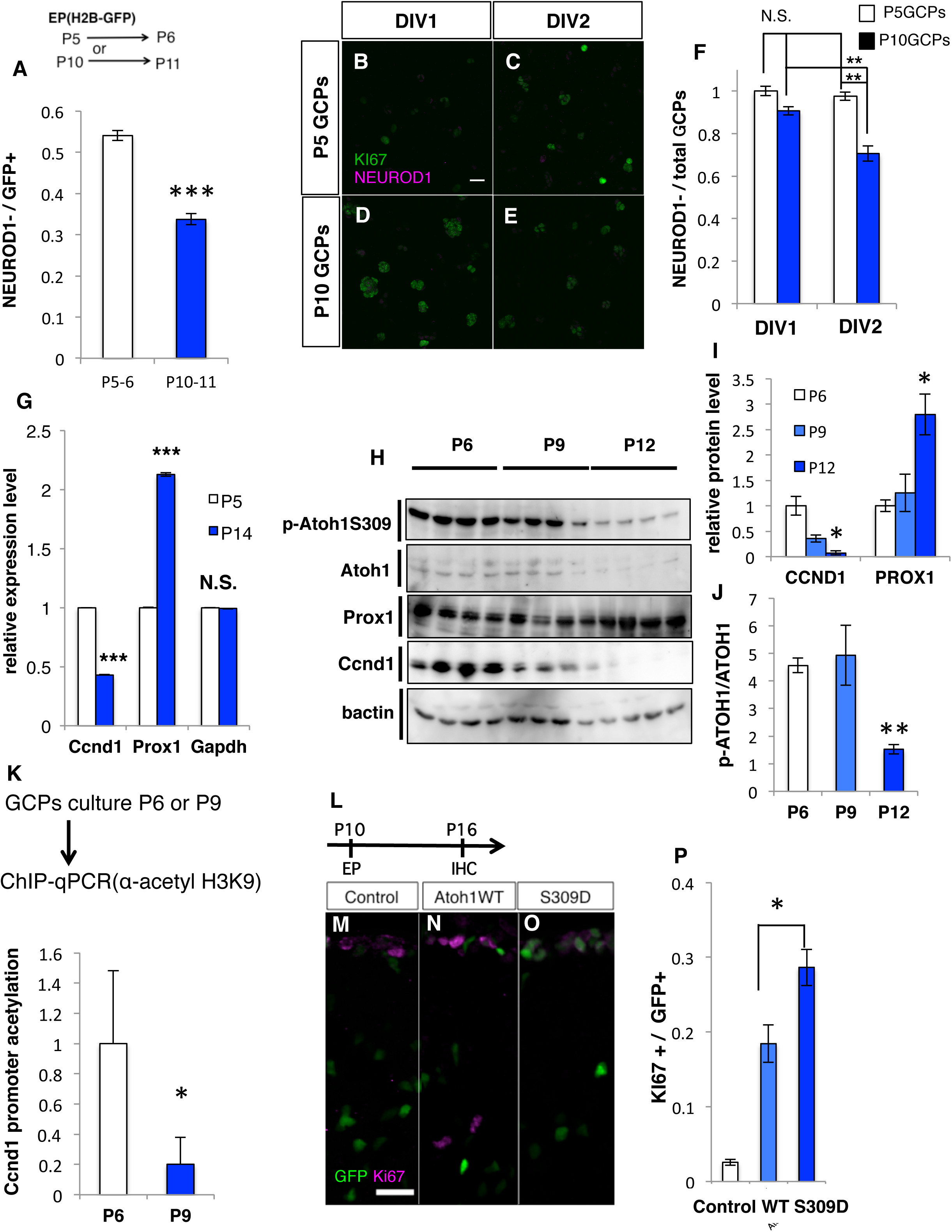
Dynamic expression of ATOH1,CCND1 and PROX1. (A) P5 or P10 mouse cerebella were electroporated with H2B-GFP plasmids and immunostained with anti-Neurod1 and anti-Ki67 the following day (P6 or P11). The ratio of NEUROD1-negative KI67-positive cells (B-F) AT-ND transition in cultured GCPs. (B-E) GCPs were extracted from P5 (P5 GCPs) or P10 (P10 GCPs) mouse cerebella and immunostained with anti-Neurod1 (magenta) and anti-Ki67 (green) at DIV1 and DIV2. (F) Ratios of NEUROD1-cells (AT+GCPs) among total KI67-positive cells. (G) Normalized levels of Ccnd1, Prox1 and Gapdh transcripts were compared between Atoh1-positive cells extracted from the scRNA-seq data of P5 and P14 cerebellum (see also Experimental Procedures). Wilcox test between n=1602 cells (P5 Atoh1-positive cells) and n=274 cells (P10 Atoh1-positive cells). (H-J) Whole cerebellum lysates of P6, P9 and P12 mice were subjected to immunoblotting with anti-pAtoh1 S309, Atoh1, Ccnd1, Prox1, β-actin. (I) Quantification of p-ATOH1 S309 normalized to the total level of ATOH1. (J) The protein levels of CCND1 and PROX1 normalized by β-ACTIN. (K) ChIP assay for H3K9 histone-acetylation of *Ccnd1* promoter in cultured GCPs extracted from P6 or P9 mouse cerebellum. Student T test between P6 GCPs (N=4) and P9 GCPs (N=4). (L-P) Over-expression analysis of WT or S309D form of Atoh1. (L) Expression plasmids of either WT or S309D form of Atoh1 were introduced to P10 mice EGL and immunostaining for KI67 (magenta) was performed in P16 mouse cerebellum. Nuclei of electroporated cells were detected by GFP (green). Images of H2B-GFP (M, Control), WT(N) and S309D (O). (P) Ratios of KI67+ cells among the electroporated cells. Tukey-Kramer test among Control (N=3), Atoh1 WT (N=3) and Atoh1 S309D (N=3). Error bar sem. Scale bars: 20um(B-E, M-O)

Analysis of scRNA-seq data for P5 and P14 (GSM3318007) cerebella revealed that *Ccnd1* expression was higher at P5 and lower at P14 in AT+GCPs, while *Prox1* expression was lower at P5 and higher at P14 in those cells (Fig. 6G). Our western analyses of whole lysates from developing cerebella revealed a similar tendency as to expression of CCND1 and PROX1 during development (Fig. 6H, I). Interestingly, the ratio of the p-S309-ATOH1 over total ATOH1 was significantly decreased at P12, although the total amount of ATOH1 protein gradually decreased during development (Fig. 6J). Furthermore, our ChIP assay using anti-acetyl H3K9 in the cerebella revealed that histone acetylation around the *Ccnd1* promoter at P9 was much lower than that at P6 (Fig. 6K). These findings suggest the following hypothesis: At early cerebellar developmental stages, PROX1 expression is weaker in AT+GCPs. In contrast, Ccnd1 expression is higher because its promoter chromatin region is less acetylated. High expression of CCND1 cooperates with CDK4 to stabilize ATOH1 by phosphorylating S309, which eventually suppresses AT-ND transition. At later stages, higher expression of PROX1 in AT+GCPs elicits chromatin deacetylation at the *Ccnd1* promoter region and reduces CCND1 expression. Reduction of CCND1 expression causes less phosphorylation of S309 ATOH1, which de-stabilizes ATOH1 and leads to AT-ND transition. During development, the gradual increase of PROX1 causes a gradual decrease of CCND1, leading to a gradual decrease in ATOH1 through phosphorylation. The gradual decrease of CCND1 and ATOH1 may account for gradual reduction of proliferation and immaturity of GCPs, respectively. While electroporation of WT-ATOH1 to P10 EGL resulted in significant increase of KI67-positive cells (putatively GCPs) at P16, such an effect was largely increased with S309D-ATOH1 (Fig. 6L-P), supporting our hypothesis.

### Canonical WNT signaling controls the dynamic expressions of CCND1 and ATOH1 through the induction of PROX1

We speculate that some molecules and/or signaling pathway might mediate the dynamic expression of PROX1 and CCND1, which would affect the protein stability of ATOH1 and subsequent AT-ND transition. Canonical WNT signaling is reported to suppress proliferation and promote differentiation of GCPs (Lorenz et al., 2011; Pei et al., 2012). In addition, PROX1 expression is upregulated by canonical WNT signaling in the cerebral cortex (Karalay et al., 2011), assuming that WNT signaling might drive PROX1 expression in the EGL. We first performed immunostaining with non-phosphorylated form of β-catenin (NP- β-catenin), whose nuclear localization indicated WNT signaling activities, in developing cerebella at P5. WNT signaling was stronger in the deeper side of EGL but very faint in the upper side (Fig. 7A-C), similar to PROX1 expression. We then electroporated the constitutive active form of β-catenin (β-cat S33Y) to P5 cerebella. One day after electroporation (P6), the ratio of PROX1-positive cells in electroporated cells was significantly increased by β-cat S33Y introduction (Fig. 7D-F), supporting that WNT signaling increases PROX1 expression in the EGL.

**Figure 7.**
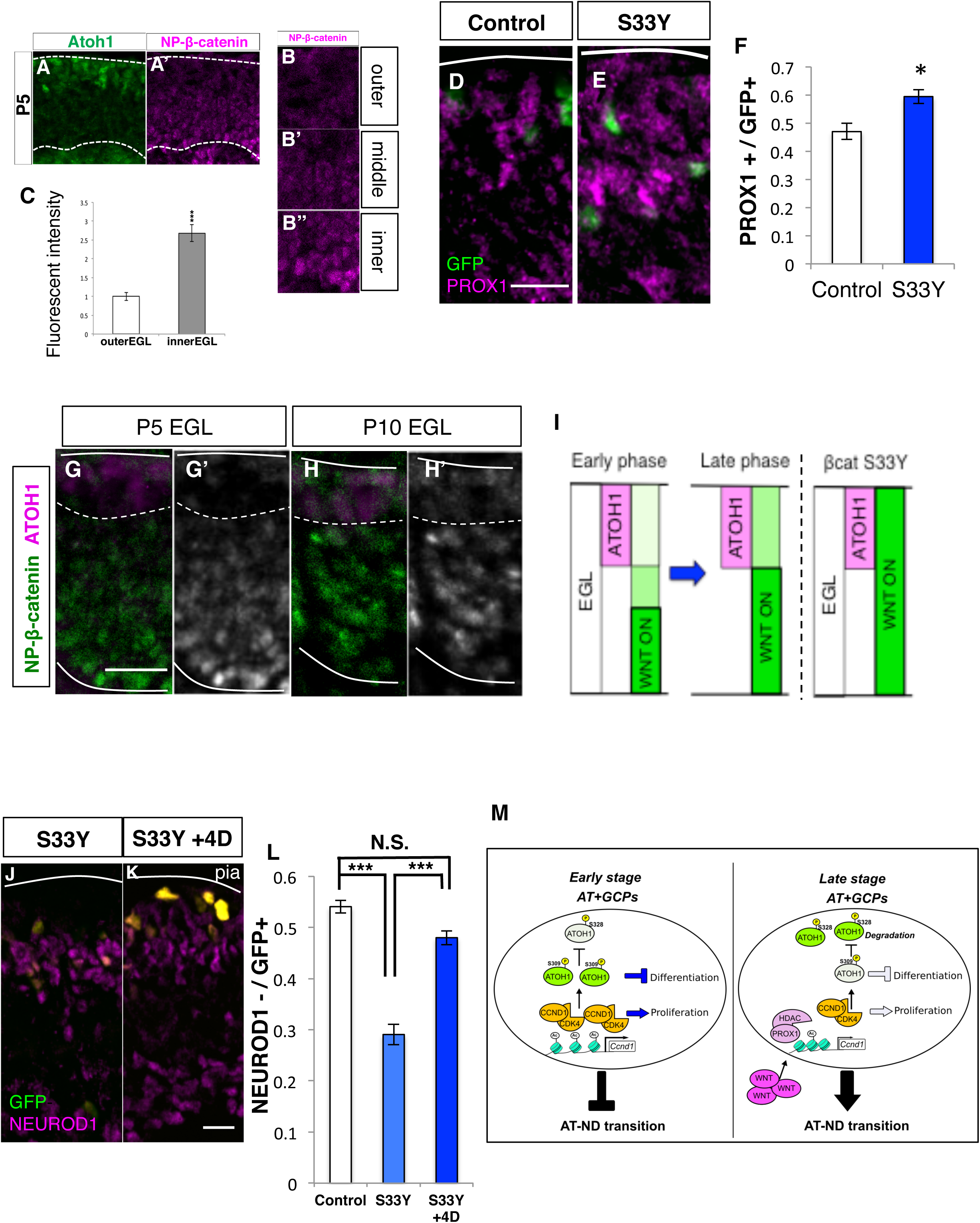
Canonical WNT signaling controls the AT-ND transition. (A) Immunostaining of P5 cerebellum with indicated antibodies. (B-C) Quantification of the intensity of NP- β-catenin in the outer and the inner parts of EGL. (D-F) Over-expression analysis of the activated form of β-catenin (βcat S33Y). (D, E) Expression plasmids of βcat S33Y were introduced to P5 mice EGL and immunostaining for PROX1 (magenta) was performed the following day. Nuclei of electroporated cells were detected with GFP (green). Images of H2B-GFP (D, Control) and βcat S33Y). (F) Ratios of PROX1-positive cells in electroporated cells. (J-L) Rescue experiment for βcat S33Y. (G-H) Immunostaining of P5 or P10 cerebellum with indicated antibodies. (I) Schematic illustrations of dynamic activity of WNT signaling estimated by the intensity of NP- β-catenin. (J, K) Expression plasmids of βcat S33Y were introduced with or without Ccnd1 and Cdk4 expression plasmids to P5 mice EGL and immunostaining for NEUROD1 (magenta) was performed the following day. Nuclei of electroporated cells were detected by GFP (green). Images of βcat S33Y (J, S33Y) and βcat S33Y with Ccnd1 and Cdk4 (K). (L) Ratios of NEUROD1 negative and KI67+ cells (AT+GCPs) among the electroporated cells. (M) Schematic illustrations summarizing this study. Scale bars: 20um(D-E, G-H, J-K)

Intriguingly, immunostaining with NP- β-catenin showed that activity of WNT signaling was increased from P5 to P10 (Fig. 7G-I), consistent with previous reports that expression of WNT ligands were increased during cerebellar development (Anne et al., 2013). Next, we electroporated β-cat S33Y to increase WNT signaling in GCPs, which resulted in significant decrease of AT+GCPs (NEUROD1-negative cells) (Fig. 7J-L). However, this phenomenon was rescued by co-introduction of CCND1/CDK4, confirming that CCND1/CDK4 antagonizes AT-ND transition downstream of WNT signaling. Taken together, the gradual increase of WNT signaling during cerebellar development may account for gradual loss of proliferation activity and immaturity through controlling PROX1, CCND1 and ATOH1 (Fig. 7M).

## Discussion

During normal development, it is important for progenitors to properly proliferate and then appropriately differentiate into mature cells. In general, progenitors in the immature state tend to proliferate actively, while mature cells, in many cases, divide slowly or cease cell division. However, quiescent stem cells such as hematopoietic stem cells, spermatogonia, and adult neural stem cells remain in immature states but do not divide rapidly (de Rooij, 2017; Homem et al., 2015; Nakamura-Ishizu et al., 2014). This suggests that proliferation and immatureness can be regulated distinctly, although they can cooperatively affect each other. This notion is supported by the phenotype of certain KO mice. In Ink4d and Kip1 double KO mice, neurons, which are supposedly differentiated postmitotic cells, actively proliferate even after completion of their migration (Zindy et al., 1999). In the telencephalon-specific retinoblastoma (Rb) KO mice, some neurons outside of the VZ were still found to divide (Ferguson et al., 2002). In the cerebellum, we reported that ATOH1-positive immature GCPs migrate inward with low proliferative activity in Meis1 KO mice (Owa et al., 2018). However, the molecular machinery to distinctly but cooperatively regulate proliferation and immatureness of progenitors remains unclear.

In this study, we found that there are two types of GCPs in the EGL, AT+GCPs and ND+GCPs, which are localized in the pial and middle region of the EGL, respectively. AT+GCPs are more proliferative and immature, and gradually give rise to the less proliferative and more differentiated ND+GCPs. Therefore, investigation of the AT-ND transition should give insight into the cooperative regulation underlying the proliferation and immatureness of progenitors. Previously, other groups have suggested the existence of subpopulations of GCPs in the EGL (Shiraishi et al., 2019; Xenaki et al., 2011). Xenaki et al reported that TAG1/CNTN2 is expressed in the middle region of the EGL, where ATOH1 is not expressed. By analyzing the scRNA-seq data, we found that Cntn2 is specifically expressed in ND+GCPs but not in AT+GCPs (Fig. S1C), suggesting that TAG1/CNTN2-positive CGPs may largely correspond to ND+GCPs.

A well-known function of CCND1 is to promote the G1-S transition of proliferating cells in cooperation with CDK proteins (Sherr, 1994). In the cerebellum, CCND1 has been reported to upregulate the proliferation of GCPs (Pogoriler et al., 2006). In contrast, ATOH1 is known to maintain GCPs in the immature state (Ayrault et al., 2010; Flora et al., 2009). However, it was unclear how CCND1/CDKs and ATOH1 reciprocally affect one another to cooperatively regulate the proliferation and immaturity states of GCPs. We found that CCND1/CDK4 stabilizes ATOH1 protein via direct phosphorylation of S309 ATOH1, which allows CCND1 not only to directly control cell cycle progression but also to indirectly regulate the immaturity of GCPs. Similar what we demonstrate for CCND1/CDK4 and ATOH1, CCND/CDK complexes have been shown to phosphorylate not only cell cycle-related proteins but also other proteins, such as transcription factors. For example, the CyclinA/CDK2 complex phosphorylates Neurogenin2, a bHLH transcription factor, thereby modulating its transcriptional activity of (Ali et al., 2011).

Many studies have suggested that phosphorylation of ATOH1 affects its protein stability (Aragaki et al., 2008; Cheng et al., 2016; Forget et al., 2014; Klisch et al., 2017; Tomic et al., 2018; Tsuchiya et al., 2007; Xie et al., 2017). Among previously reported phosphorylation sites, S52, S56, tyrosine (Y)78, S193, S328, S334 and S339 of mouse ATOH1 are believed to participate in the protein stability. Tsuchiya et al showed that GSK3β phosphorylates S54 and S58 (corresponding to S52 and S56 in mouse) in a human colon cancer cell line, which promotes ATOH1 degradation. Forget et al reported that phosphorylation of S328 and S339 of mouse ATOH1 leads to proteasome-dependent ATOH1 degradation by the E3 ubiquitin ligase, HUWE1. Cheng et al also showed that Casein kinase 1 phosphorylates S334 of mouse ATOH1 to reduce the ATOH1 stability via the action of HUWE1. Xie et al further showed that phosphorylation at S193 of mouse ATOH1 increases ATOH1 degradation in HEK293T cells, although they did not identify the responsible protein kinase. In knock-in mice where S193 is substituted with an alanine residue, hair cells in the cochlea undergo degeneration, suggesting the importance of ATOH1 S193 phosphorylation in inner ear development. Moreover, Klisch et al highlighted that JNK2 phosphorylates Y78 of ATOH1, which stabilizes ATOH1. While this tyrosine phosphorylation was only weakly observed in the normal cerebellar tissue, elevated phosphorylation of Y78 was found in medulloblastoma. Reduction of this phosphorylation decreased proliferation of medulloblastoma cell lines. Tomic et al generated knock-in mice that expressed a mutant ATOH1 protein with alanine substitutions for nine proline-directed serine/threonine residues. These substitutions resulted in a complete block of ATOH1 phosphorylation and reduction of the pool of stem cell progenitors in the intestinal epithelium, lowering its regeneration potential. These findings suggested the importance of phosphorylation in ATOH1 stability and in a variety of biological events, such as inner ear development, intestinal epithelium regeneration and medulloblastoma growth.

Although it was reported that S309 of ATOH1 can be phosphorylated in HEK293T cells, the responsible protein kinase as well as the effect on the protein stability were unknown. In this study, we found that CCND1 and CDK4 cooperatively phosphorylate S309 of ATOH1, resulting in stabilization of ATOH1 protein in N2a cells. Moreover, our experiments using specific antibodies against pS309-ATOH1 and pS328-ATOH1 revealed a unidirectional relationship between the two phosphorylation sites. If S309 of ATOH1 is phosphorylated, S328 cannot be phosphorylated. However, phosphorylation of S328 does not inhibit further phosphorylation at S309. Because S328 phosphorylation causes ATOH1 degradation via the action of the ubiquitin E3 ligase, HUWE1, phosphorylation at S309 by CCND1/CDK4 may suppress ATOH1 degradation via inhibition of S328 phosphorylation. This is the first report on such crosstalk among the phosphorylation sites of ATOH1 and, we believe, provides good insights into the machinery to control protein stability by multiple phosphorylation.

WNT signaling is one of the most fundamental signals that control tissue development (Steinhart and Angers, 2018). In the developing cerebellum, canonical WNT signaling is known to promote differentiation from GCPs to GCs (Lorenz et al., 2011; Pei et al., 2012). Although the molecular machinery underlying how canonical WNT signaling accelerates the GC differentiation is still not well understood, our experiment with β-catenin S33Y overexpression revealed that WNT signaling promotes AT-ND transition by upregulating PROX1. WNT7a is expressed in GCs and thought to be the ligand for canonical WNT signaling in GCPs (Anne et al., 2013; Lucas and Salinas, 1997). As development proceeds, WNT7a expression is increased in the cerebellum. In this study, we revealed that canonical WNT signaling upregulates PROX1 expression, that PROX1 downregulates CCND1 expression, and that CCND1 in cooperation with CDK4 phosphorylates and stabilizes ATOH1. Therefore, a gradual increase in WNT7a expression may account for gradual increases in canonical WNT signaling and PROX1 protein levels as well as gradual decreases of ATOH1, p-S309-ATOH1 and CCND1, which together may contribute to maintaining the appropriate AT-ND transition at specific developmental stages. SHH signaling is another well-known extrinsic cue that maintains GCPs in immature and proliferative states (Wallace, 1999). In the cerebellum, Purkinje cells express and secrete SHH, which gradually decreases as development proceeds. Thus, the antagonistic environmental signals of SHH and WNT may coordinately regulate the balance of GCP proliferation and differentiation, although the mechanisms for such crosstalk between these two signals is still elusive.

In the telencephalon of mammals, including humans, there is a transient or secondary mitotic zone containing intermediate progenitors and/or outer radial glia that contribute to expansion of the brain (Arai and Taverna, 2017). Acquisition of these secondary progenitors during evolution is thought to contribute to the development of a larger and more complex cerebral cortex (Florio and Huttner, 2014). The cerebellum has been getting bigger during vertebrate evolution. In Zebrafish and Xenopus, postmitotic GCs are directly produced from the rhombic lip, although Xenopus has an EGL-like structure that does not contain mitotic GCPs (Butts et al., 2014a). Avians have acquired the secondary amplifying cells, GCPs, in the EGL, which contributes to the expansion of the size and complexity of the cerebellum (Butts et al., 2014b). In this study, we further found that mice have both secondary and tertiary amplifying cells, AT+GCPs and ND+GCPs, respectively. Because avians do not have ND+GCPs (Butts et al., 2014b), it suggests that the addition of this third amplifying system enables mammals to further expand their cerebellar structure. Additional studies will be required to evaluate how proliferation of ND+GCPs affects the evolutionary expansion of the cerebellum.

Pediatric tumors, medulloblastomas (MBs), can be classified into four subgroups (WNT-type, SHH-type, Group 3, Group 4) based on clinical and molecular characteristics (Northcott et al., 2012). SHH-MBs are known to emerge from the cerebellar granule cell lineage. A recent comprehensive scRNA-seq study showed that infant SHH-MB and adult SHH-MB have different molecular characteristics. While adult SHH-MBs express Atoh1, infant SHH-MBs express genes such as *Neurod1*, *Map1b* and *Tubb2b* (Hovestadt et al., 2019), which are strongly expressed in ND+GCPs (Fig. 1e). We suspect that ND+GCPs may be the origin for infant SHH-MBs, although further studies would be required.

The proliferative activity and immature state of progenitor cells are not necessarily synchronized, but they are often coupled together during tissue development. In this study, we revealed that CCND1 coordinately controls proliferation activity and immaturity in GCPs during cerebellar development. CCND1 not only promotes cell cycle progression but also contributes to the maintenance of immaturity by stabilizing ATOH1. Our results contribute to understanding the machinery for proliferation and differentiation of cells in many tissues and provide insights into the pathogenesis of MBs.

## Acknowledgement

We would like to thank Dr. Ruth Yu for grateful comments. This work is supported by Grants-in-Aid for Scientific Research (Grant 18H02538 to M.H. and 15J06259 to S.M.) and Innovative Areas (Grant 16H06528 to M.H.) from MEXT, Strategic Research Program for Brain Sciences from AMED (JP16dm0107085h0001), Takeda Foundation and Intramural Research Grants (Grants 30-9 and 1-4 to M.H.).

## Author Contributions

Conceptualization, S.M., Y.S. and M.H; Methodology, S.M. T.O. and Y.S.; Software, S. M.; Varidation, S.M.; Investigation, S.M., T.O., Y.S and T.N.; Resources, S.M., T.O., Y.S., M.Y., S.A., Y.K. and S.T.; Writing – Original Draft, S.M. and M.H.; Writing – Review & Editing, M.H.; Visualization, S.M.; Supervision, Y.S.;Funding Acquisition, S.M., S.T. and M.H.

## Declarations of interest

The authors declare no competing interests.

## Experimental Procedures

### Animals

All mouse experiments were approved by the Animal Care and Use Committee of the National Institute of Neuroscience, Japan (Approval number 31-19). Mice were housed in SPF with food and water ad libitum. ICR pups were obtained from SLC (Japan). Prox1-floxed mice were a generous gift from Dr. Y. Kawaguchi (Goto et al., 2017). For immunohistochemistry, neonatal mice were fixed with 4% PFA and embedded sagittally with O.C.T compound (Sakura Finetek). Frozen brains were sectioned into thin slices via cryostat (Leica Biosystems). For immunoblotting, neonatal mouse cerebellum was removed and mechanically triturated in ice-cold 1xPBS with protease inhibitor cocktail. Tissues were then placed into lysis buffer (1% NP40, 1mM NaF) and subjected to sonication. Finally, SDS sample buffer was added to lysate supernatants.

### Plasmids

The expression vectors of Atoh1, Ccnd1, Prox1, Cdk4 and βcatenin were cloned from cDNAs from C57/BL6 mouse cerebellum. Cloned fragments were inserted into a pCAGGS vector or pGEX-4T vector (GE Healthcare) and cloned sequences confirmed by sequencing. pCAG-Cre vectors were cloned from CAG-Cre transgenic mice (Hori et al., 2014). pCAG-H2BGFP vectors were a gift from Dr. N. Masuyama. Mutated forms of Atoh1 S309A, S309D and βcatenin S33Y were generated following the mutagenesis protocol of PrimeStarMax (Takara). shRNAs were generated by inserting the double-stranded oligonucleotides into a mU6 pro vector. The targeting sequence of each shRNAs was designed by siDirect 2.0 (Naito et al., 2004; Naito et al., 2009) and sequences are indicated below.

### Cell culture, transfection and drug treatment

Neuro2a (N2a) cells were obtained from ATCC and cultured in DMEM containing 10%FBS and 100U/mL Penicillin-Streptomycin. Transfection of N2a cells was performed with Transfectin reagent (Bio-Rad). 100uM Cycloheximide (CHX) was diluted in DMSO and added to N2a cells as indicated. 100nM Palbociclib (Palb) diluted in MilliQ (excepted for the Fig 3G) and 10uM MG132 diluted in DMSO were administered to N2a cells. Cells were harvested at indicated times.

### Immunohistochemistry

Detailed protocols were described previously (Seto et al., 2014). In brief, cryosections were incubated at room temperature with 1% normal donkey serum containing 0.2% PBST (blocking solution) for one hour. After blocking solutions were removed, antibodies listed in KEY RESOURCE TABLE were diluted in blocking solution and applied to sections. Each section was incubated at 4°C for 16 hrs. Donkey-raised secondary antibodies (abcam) were then applied onto sections. Cryosections were incubated for 2 hours at room temperature. After incubation, DAPI staining was performed to visualize nuclei and each section was mounted with Permaflour.

### Cumulative Labeling Method

50mg/kg EdU was injected intraperitoneally into P6 ICR mice every 3 hours to perform cumulative labeling. Mice were fixed with 4% PFA 30 min after EdU injection. EdU+ cells were detected with the EdU Click-iT kit (Thermo) following the manufacturer’s protocol. Coimmunostaining was performed for Atoh1 and Ki67. To determine the cell cycle length of each progenitor, the ratios of EdU+ cells among the ATOH1+ KI67+ cells (AT+GCPs) or ATOH1-KI67+ cells (ND+GCPs) located around the central EGL were measured with ImageJ from N=3 mice per each time point. Cumulating curves were calculated as previously described (Takahashi et al., 1995) based on the 0, 3, 6 and 9 hr data.

### in vivo Electroporation

in vivo electroporation in neonatal mice was described previously (Owa et al., 2018). Expression plasmids were diluted to 1 ug/ul, shRNAs to 2 ug/ul and pCAG-H2BGFP to 0.25ug/ul in milliQ. Fastgreen was added to visualize the plasmid solution. Plasmid solutions were injected into P5 or P10 ICR cerebella over the skull. 50msec length 80V electric pulses were delivered to mice with 150msec interval using forceps-type electrodes (NEPA gene). The pups were kept warm at 37°C to recuperate and returned to the litter after fully recovering. Pups were fixed with 4% PFA 1 day or a few days after electroporation. To inhibit HDAC activity, 50mg/kg Valproic acid (VPA) was administered intraperitoneally to P6 and P7 pups followed by electroporation.

### Immunoblotting

Detailed protocols were described previously (Shiraishi et al., 2019). Briefly, transferred PVDF membranes were incubated with primary antibodies listed on the KEY RESOURCE TABLE at 4 degrees, overnight. After membranes were incubated with secondary antibodies at RT for 2 hr, HRP substrate (Millipore) was applied and immuno-signals were detected by LAS4000 (Fujifilm).

### GST-pull down, Protein purification and Phospho-mass spectrometry

GST-tagged Atoh1, Atoh1S309A, Ccnd1 and Cdk4 were transfected to N2a cell lines and cultured for 2 days. N2a cells were harvested as described above. After lysis, supernatants were incubated with glutathione conjugated sepharose-beads at 4°C overnight. Sepharose-beads were washed 2 times and resuspended with sample buffer. For protein purification, washed sepharose-beads were incubated with TED buffer containing 150mg reduced-form glutathione (pH 7.4) in 4 °C for 30 min and supernatants were stored at −80°C. For λ-phosphatase assay, purified proteins were incubated with λ-phosphatase at 30°C for 30min. After that, sample buffer was added to the solution containing the purified protein. Controls were handled with same protocol without λ-phosphatase (NEB). Phospho-mass spectrometry was performed as previously described (Nagai et al., 2016).

### In vitro kinase assay

3-5ug purified ATOH1 proteins were incubated with TED buffer containing 20uM ATP, 0.5uCi [γ-32P]-ATP, 20 mM MgCl2 and 3-5ug kinase (CK1α or CCND1/ CDK4) at 37°C for 1 hour. Reacted solution was then applied to sample buffer and subjected to SDS-PAGE. Phosphorylation signals were detected by Typhoon (GE healthcare).

### ChIP assay

ChIP assay was performed as previously described. Briefly, P5 mouse cerebella were removed and cross-linked by 1% formic acid. Tissues were sonicated in the lysis buffer (1%SDS, 10mM EDTA, 50mM Tris-Hcl) containing the proteasome-inhibitor cocktail and diluted to 1/10 with the dilution buffer (16.7 mM Tris-Hcl, 167 mM NaCl, 1.1% TritonX 100, 1.2 mM EDTA, 0.01% SDS), which was used as an input. 50-150 ug of DNA fragments were incubated with 2ug Prox1 antibody (control: 2ug goat IgG) and protein A sepharose beads at 4°C overnight. The following day, DNA fragments bound to antibodies were washed and eluted in the elution buffer (1%SDS, 0.1 M NaHCO3). DNA fragments were then reverse-cross-linked with 5M NaCl at 65°C for 4 hours. DNA fragments were incubated with Proteinase K and RNase in 45°C for 1 hour and extracted by phenol-chloroform. DNA fragments were amplified in the Ccnd1 promoter region by PCR with the following primers.

### Cerebellar primary culture

P5 or P10 mouse cerebella were extracted and meninges and choroid plexi were removed in HBSS. The tissues were dispersed with Neuron Dissociation Solutions (FujiFilm) and plated in poly-D-lysine coated dishes. Primary cells were cultured in DMEM supplied with 1xN2, 1xB27, 10% FBS, 1uM SAG and 100U/mL Penicillin-Streptomycin. For immunocytochemistry of primary cerebellar culture, cultured cells were fixed with 4% PFA for 20 min at RT and 10% normal donkey serum diluted with 1% PBST was applied to cultured cells. Cultured cells were incubated with Neurod1, Pax6 and Ki67 antibodies diluted in blocking solution at 4°C overnight.

Secondary antibodies were added the following day after cultured cells were washed with PBS for 2 hours in RT. After incubation, DAPI staining was performed to visualize nuclei. For ChIP assay, 1×106 cells were subjected to ChIP assay following the above protocol. Acetyl-H3K9 antibody was used for precipitation and primer sets were same as above.

### Data processing for published single cell RNAseq data

All data sets used in this study were obtained from GEO(accession number) and analyzed with Seurat v3 (Stuart et al., 2019) according to the author’s protocol. In brief, prospective GCPs (Pax6>1 &&Pcna >1) were extracted after pre-processing and normalizing the original data sets. Cell cycle score of each GCP was calculated with default settings and cell clustering (resolution=0.1) was performed with regressing out the effect of cell-cycle (Tirosh et al., 2016). UMAP based dimensional reduction, differentially expressed features and heatmap images were obtained with Seurat.

### Image acquisition and quantification

All images were acquired at midline vermis with confocal microscopy, LSM780 (Carl Zeiss) and FV3000 (Olympus). Acquired images were analyzed with Image J (RSB).

### Statistical analysis

Individual animals or trails are regarded as biological replicates. All data are presented as mean ± S.E.M., also indicated in the each Figure legend. Statistical tests were performed by Student’s t test or Tukey’s test. p-values were represented as; N.S. for p>0.05; *, p<0.05; **, p<0.01; ***, p<0.001. The same controls were used for statistical test under the same experimental conditions.

### Code availability

All the codes used in this study are available from the lead contract.

## Supplemental Information titles and legends

**Figure S1 Related to** **Figure 1**

(A) Immunostaining with indicated antibodies to P6 mice. 50mg/kg EdU was administered intraperitoneally 30 min before fixation. Solid white arrowheads indicate NEUROD1-positive and EdU-positive GCPs (ND+GCPs in S-phase). White hollow arrowheads indicate NEUROD1-negative and EdU-positive GCPs (AT+GCPs in S-phase). (B) Immunostaining with indicated antibodies to P6 mice. Solid white arrowheads indicate NEUROD1-positive and PH3-positive GCPs (ND+GCPs in M-phase). White hollow arrowheads indicate ATOH1-positive and EdU-positive GCPs (AT+GCPs in M-phase). (C) Normalized expression of selected genes (Pax6, Pcna, Plxnb2, Barhl1, Mki67, Mcm6, Ccnd1 and Cntn2) were mapped onto the UMAP dimension. (D, E) Efficiency of shRNAs was confirmed by immunoblotting. Lysates of N2a cells transfected with either Atoh1 expression plasmids with scramble (Scr) shRNA plasmids or Atoh1 expression plasmids with Atoh1 shRNA (sh#1, sh#2 or sh#3) were subjected to immunoblotting with anti Atoh1 or β-actin (D). Lysates of N2a cells transfected with either GFP expression plasmids fused to Atoh1 3’ untranslated region (UTR) (GFP-3’UTR) with scramble (Scr) shRNA plasmids or GFP-3’UTR with Atoh1 sh#3 targeting 3’UTR region of Atoh1 were subjected to immunoblotting with anti GFP or β-actin (E). Scale bars: 20um(A-B)

**Figure S2 Related to** **Figure 2**

(A) Cell cycle-related expression of Ccnd1 in GCPs categorized to cluster 0 (AT+GCPs) or cluster 1 (ND+GCPs, Fig 1C). (B) Immunostaining with indicated antibodies in P6 mice. Strong CCND1 expression was observed in dividing AT+GCPs (NEUROD1-negative and PH3-positive), while CCND1 expression was faint in ND+GCPs (NEUROD1-positive and PH3-positive). (C) Efficiency of shRNAs was confirmed by immunoblotting. Lysates of N2a cells transfected with either sh-scramble plasmid (scr) or Ccnd1 shRNA (sh#1 or sh#2) were subjected to immunoblotting with β-actin to examine endogenous expression of CCND1. (E-G) Ccnd1 KD analysis related to Fig. 2C-F. (E, F) Ccnd1 shRNAs were introduced to P5 EGL with H2B-EGFP expressing plasmid and immunostaining for Neurod1 (blue) and Ki67 were performed the following day. Nuclei of electroporated cells were detected with GFP (green). Images of Control (E) and Ccnd1 KD (F). Scale bar 20um. (G) Ratios of ND+GCPs (NEUROD1-positive and KI67-positive cells) among the electroporated GCPs (GFP+, KI67+ cells). The Student T-test was performed between Control (N=3) and Ccnd1 KD (sh#1: N=3, sh#2:N=3) mice. Error bar sem. Scale bars: 10um(B), 20um (E-F).

**Figure S3 Related to** **Figure 3**

(A) GST-tagged ATOH1 or ATOH1S309A were purified from N2a cells with glutathione sepharose beads. Purified ATOH1 was incubated with λ-phosphatase (λPP) for 30 min. Purified proteins were resuspended and subjected to immunoblotting with indicated antibodies. (B) Immunoblotting with the indicated antibodies against lysates of N2a cells transfected with either Atoh1-GST or Atoh1 S309A expression plasmids.

**Figure S4 Related to** **Figure 4**

(A) GST-tagged ATOH1 was purified from N2a cells. Purified ATOH1 was incubated with λ-phosphatase (λPP). Purified proteins were resuspended and subjected to immunoblotting with indicated antibodies. (B) Immunoblotting with the indicated antibodies against lysates of N2a cells transfected with either Atoh1-GST or Atoh1 S328A expression plasmids.

**Figure S5 Related to** **Figure 5**

(A-D) Immunostaining of P6 mice with the indicated antibodies. (E) Schematic illustration of PROX1 expression in the EGL. (F) The ratio of KI67-positive cells 1 day after electroporation of Prox1 expression plasmid related to Fig. 5B-D. (G) The ratio of ND+GCPs (NEUROD1-positive and KI67-positive cells) 1 day after electroporation of Prox1 expression plasmid related to Fig. 5B-D. Scale bars: 20um (A-D).

