## Supplementary figures and images for "CyclinD1 controls development of cerebellar granule cell progenitors through phosphorylation and stabilization of ATOH1"

### Supplemental Figures

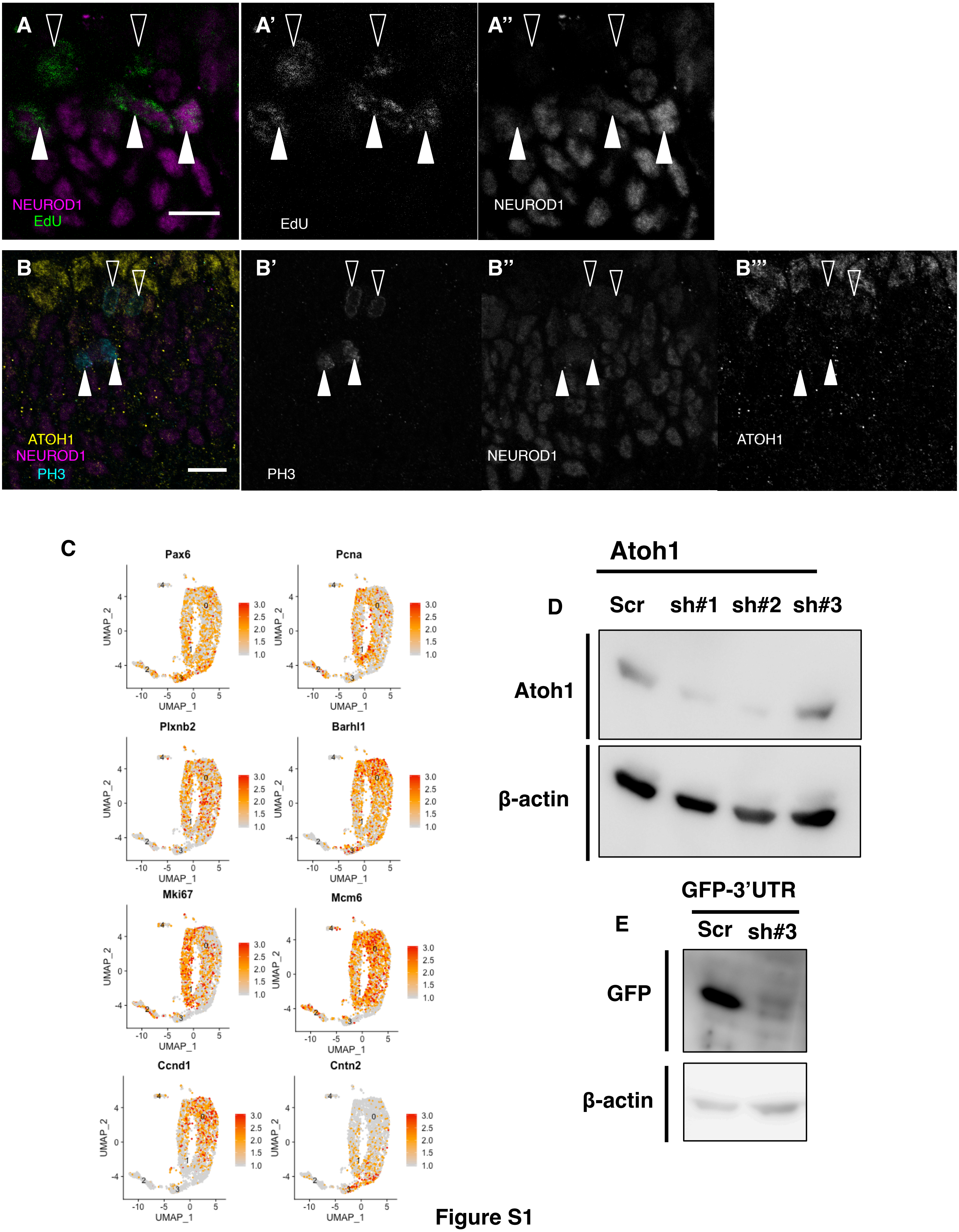

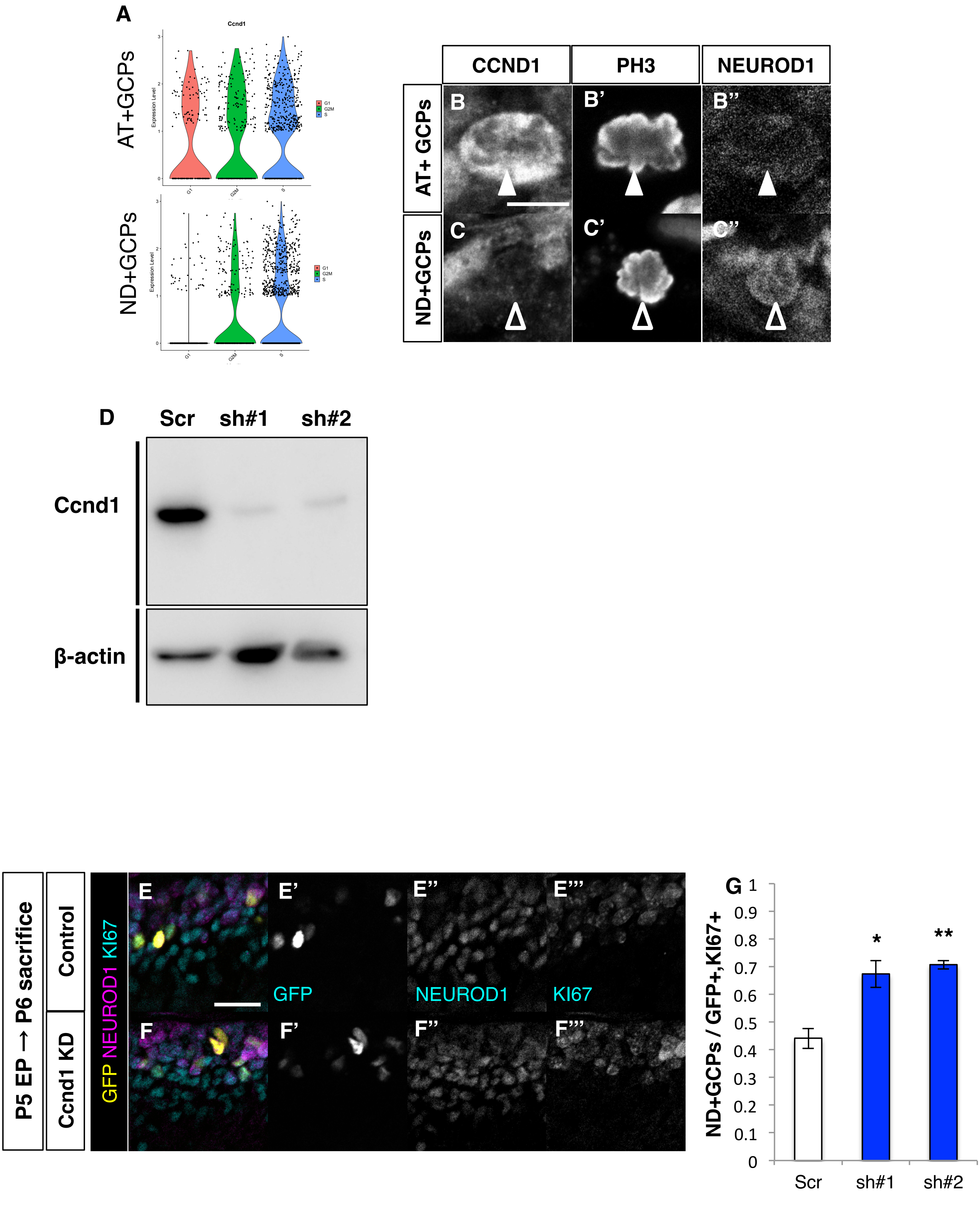

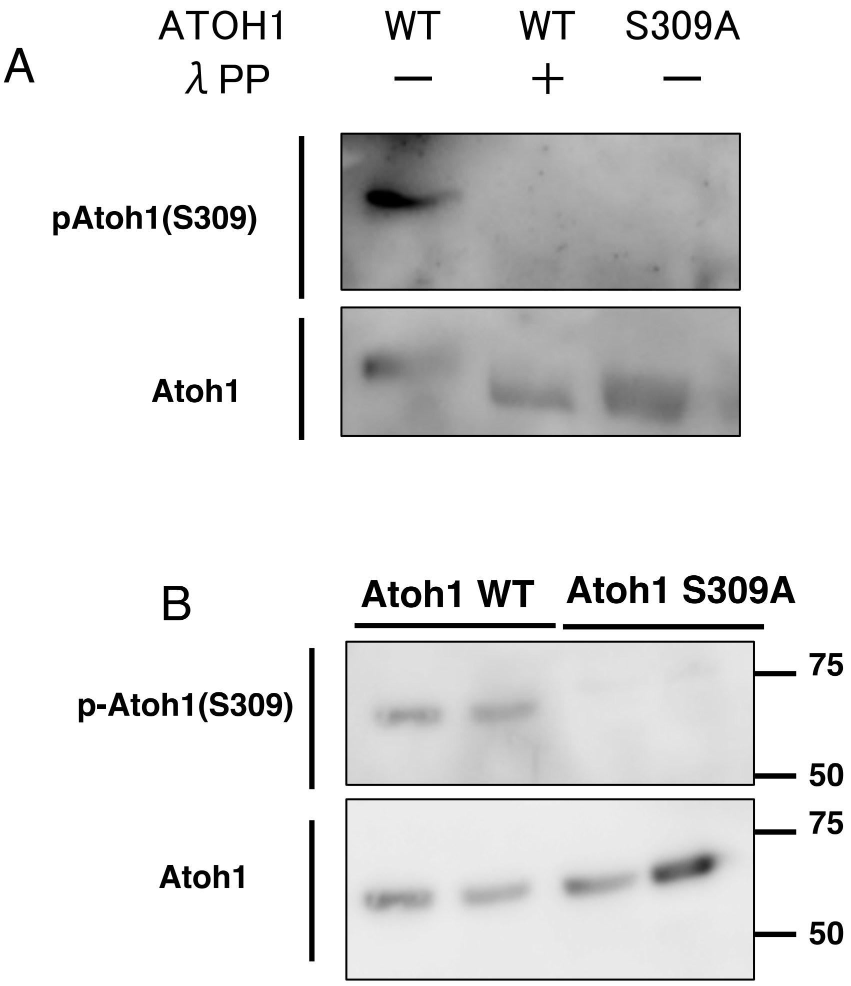

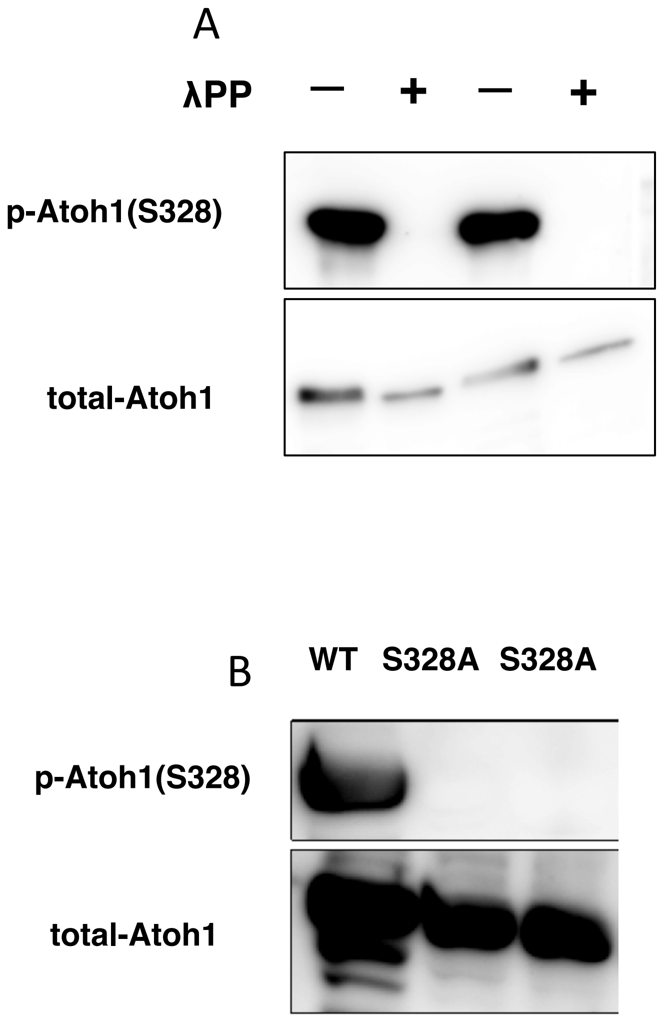

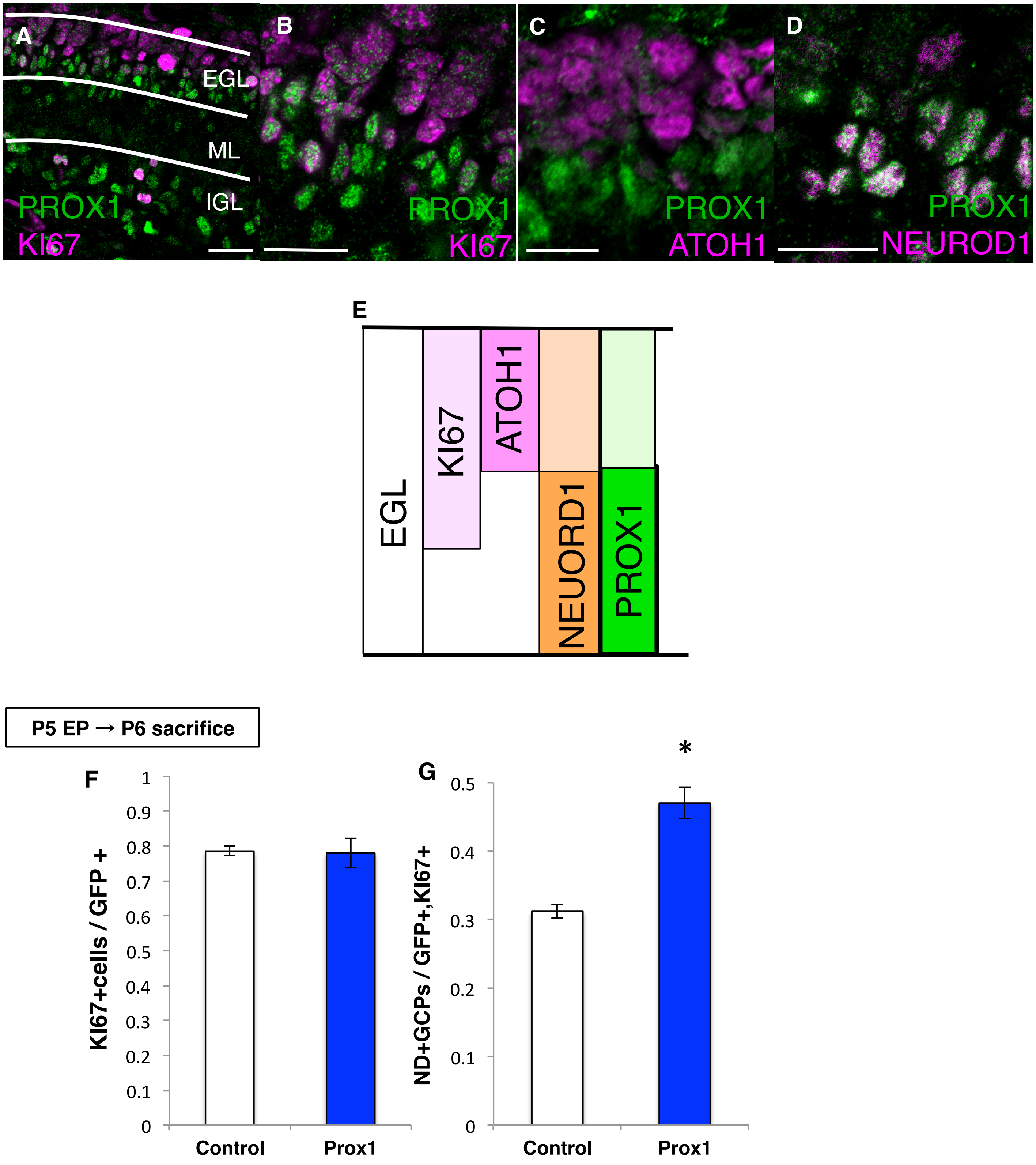
